# The Primacy of Temporal Dynamics in Driving Spatial Self-organization of Soil Redox Patterns

**DOI:** 10.1101/2023.03.28.534585

**Authors:** Xiaoli Dong, Daniel de Richter, Aaron Thompson, Junna Wang

**Affiliations:** Department of Environmental Science and Policy, University of California, Davis, California, 95616, U.S.; Nicholas School of the Environment, Duke University, Durham, North Carolina, 27708, U.S.; Department of Crop and Soil Sciences, University of Georgia, Athens, Georgia 30602, U.S.

**Keywords:** Environmental Variability, Iron Dynamics, Pattern Formation, Redox Oscillations, Soil Mottles, Turing Instability

## Abstract

In this study, we investigate mechanisms that generate regularly-spaced, iron banding in upland soils. These redoximorphic features appear in soils worldwide, but their genesis has been heretofore unresolved. Upland soils are highly redox dynamic, with significant redox fluctuations driven by rainfall, groundwater changes, or irrigation. Pattern formation in these highly dynamic systems provides an opportunity to investigate the temporal dimension of spatial self-organization, which is not often explored. By comparing multiple alternative mechanisms, we find that regular redox patterns in upland soils are formed by coupling two sets of scale-dependent feedbacks (SDF), the general framework underlying Turing instability. The first set of SDF is based on clay aggregation and disaggregation. The second set is realized by threshold-dependent, negative root responses to aggregated crystalline Fe(III). The former SDF amplifies Fe(III) aggregation and crystallinity to trigger the latter SDF. Neither set of SDF alone is sufficient to reproduce observed patterns. Redox oscillations driven by environmental variability play an indispensable role in pattern formation. Environmental variability creates a range of conditions at the same site for various processes in SDF to occur, albeit in different temporal windows of differing durations. In effect, environmental variability determines mean rates of pattern-forming processes over the timescale relevant to pattern formation and modifies the likelihood that pattern formation will occur. As such, projected climate change might significantly alter many self-organized systems, as well as the ecological consequences associated with the striking patterns they present. This temporal dimension of pattern formation is previously unreported and merits close attention.

**Statement of Significance:** Iron reactions create redox features in soils around the world. This study investigates mechanisms forming regularly-spaced iron stripes in upland soils. Upland soil redox conditions, driven by environmental variability, are highly dynamic. We show that two sets of scale-dependent feedbacks are coupled to form redox patterns and environmental variability plays a critical role in both. Significantly, environmental variability creates opportunities for various pattern-forming processes to occur at the same site in different temporal windows and determines mean process rates over the timescale relevant to pattern formation. Hence, environmental variability dictates the likelihood of pattern formation. Such a critical role of the temporal dimension in spatial self-organization has rarely been reported and has great potential for application in other self-organized ecosystems.

## 1. Introduction

Redoximorphic features are observed in soils worldwide (1, 2). During anoxic periods, reduced iron [Fe(II)] moves in soil pore water and when oxidized, forms redoximorphic features often characterized by the distinct presence (orange) or absence (gray) of Fe (3) (Fig. 1). While redoximorphic features mostly occur in soils that are frequently flooded, in drier upland soils, striking tiger-stripe-like patterns, alternating between Fe-depleted gray bands and Fe-oxide-rich orange bands, have been reported (4–6) (Fig. 1A; Table S1). The mechanism responsible for these regular patterns remains unknown; however, similar patterns are commonplace across a range of natural systems and several theories have emerged to explain them (7–10). Most notably, regular patterns in animal skin were first explained mathematically by Turing in the 1950s (11), and are thus called ‘Turing patterns’ or ‘Turing instability’. The general theory of spatial self-organization explains a broad range of patterns, e.g., vegetation patterns in drylands (12–14), regular spaced fairy circles in Namibian and Australian grasslands (10, 15, 16), patterned grounds with sorted stone and fine-grained soils in cold regions (17–19), and labyrinthine mussel beds (9, 20) and wetlands (21, 22). We investigated this a yet unexplained mechanism of redox pattern formation, a phenomenon reported in soils around the world (4–6).

**Figure 1.**
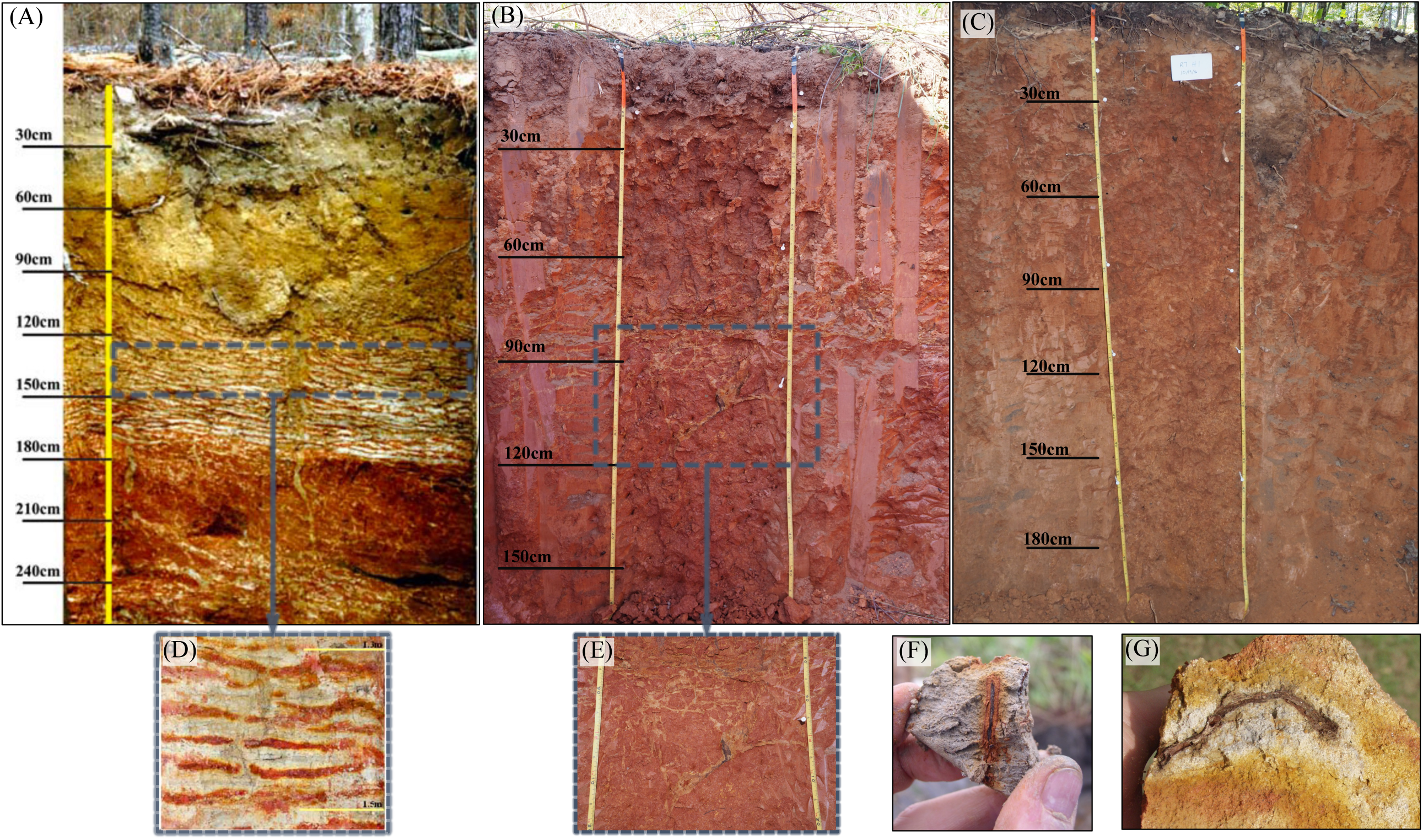
Variations of iron redox patterns in soils. (A) Our study site with extensive regular patterns, zooming into the pattern formation zone (D). (B) A site nearby (∼ 4,000 m) our study site showing narrow, irregular redox patterns, with the pattern zone zooming in (E); (C) A site nearby (∼ 400 m) our study site without any patterning; (F) rhizosphere creates an oxidized orange zone in wetland soils, while (G) rhizosphere creates a reduced gray zone in upland soils experiencing periodic inundation. Patterns shown in (A) – (E) are all from Calhoun Long-term soil experiment forests in South Carolina (U.S.).

In contrast to most of the self-organized systems studied so far, temporal dynamics of upland soils are highly complex. Upland soils undergo significant redox fluctuations driven by rainfall, groundwater table changes, or agricultural irrigation (23, 24) and redox conditions dictate soil biogeochemical processes. Environmental variability, therefore, provides opportunities for distinct processes to occur. Over the timescales relevant for pattern formation, constituent processes required for pattern formation might be able to occur at the same site, although in different temporal windows. Environmental variability further determines the duration of various processes; hence it controls the *mean rates* of pattern-forming processes over the relevant timescale. Since pattern formation requires specific configuration of relative process rates (11, 12), environmental variability may significantly alter both the occurrence and shape of patterns formed. The temporal dimension of pattern formation is thus crucial but has been heretofore rarely, if at all, investigated.

Turing instability involves a reaction-diffusion system incorporating fast-moving inhibitors and slow-moving activators, forming scale-dependent feedbacks (SDF)—short-ranged positive feedbacks and long-ranged negative feedbacks. SDF has recently been used to describe the emergence of regular vegetation patterns in drylands (12–14), where vegetation patches enhance water retention locally (short-ranged feedback), but dry out the adjacent zones (long-ranged feedback) to establish repeating spatial patterns. Soil redox oscillations appear to also provide a basis for SDF. In soils, during anoxic episodes, reductive dissolution releases micronutrients, stimulating root growth (23, 25). The resultant accumulation of organic matter (a strong reductant) from root exudates or root death reinforces reductive reactions that further release micronutrients in the rhizosphere. This forms the *short-ranged positive feedback* for root growth and expansion. The *long-ranged negative feedback* begins with Fe(II) diffusing away from the rhizosphere and, during oxic episodes, oxidizing to Fe(III). Repeated redox oscillations promote precipitation of Fe(III) of increasing crystallinity (26), which further cements into plinthite-like material (27–29). Crystalline Fe(III) weakens soil aggregation (30), which reduces soil water and nutrient retention (31, 32). Furthermore, hard and brittle plinthite-like material forms a barrier to roots (33), especially during dry periods when resistance to root penetration increases markedly (34). Together these processes form the *long-range negative feedback* that inhibit root expansion. Processes involved in the hypothesized SDF occur under different redox conditions; hence, at different times and durations during the year. For example, reductive dissolution is limited to (near) anoxic conditions, which are rare for upland soils, while root growth and expansion occur year around, subject to either stimulatory or inhibitory effect from the environment. How does such a temporal separation of processes affect pattern formation?

Functional forms of feedbacks further affect Turing instability (35, 36). For example, the effect of resources on root growth is often assumed to be monotonic—high resource concentration simulates growth while low concentration negatively affects growth; however, biological responses can also be non-monotonic. This is especially true of inhibitors, e.g., the negative effect of Fe(III) on root growth may not operate until a threshold degree of crystallinity or aggregated hardness (plinthite) is reached. A threshold dependent response requires processes that can generate and aggregate the inhibitor to trigger Turing instability (Fig. 2). Redox oscillations might further provide a mechanism to trigger the inhibitory effect of Fe(III) on root growth. Rapid pH shifts accompanying redox oscillations can enhance clay disintegration and coagulation (37). During a reducing episode, increased pH markedly disperses mineral and organic colloids (37). Soil microsites of relatively higher clay content exhibit higher water holding capacity. As a consequence, as soils dry, microsites with higher clay will be relatively wetter, with accelerated Fe(III) reductive dissolution (38). This further increases pH and enhances disintegration of silt-sized kaolinite colloids, resulting in even higher clay content. These processes form *a local positive feedback* (Fig. 2A). The large amount of Fe(II) produced in high-clay microsites diffuses to adjacent microsites and precipitates in microsites of relatively low clay (Fig. 3B), as low-clay microsites are more likely to have oxic conditions given their lower water holding capacity. In low-clay microsites, Fe(II) oxidative precipitation produces protons that lower pH and can maintain the same kaolinite colloids in silt-sized aggregates (39), decreasing the functional clay content for the already low clay microsites. This forms the *long-range negative feedback*. These processes amplify the spatial heterogeneity of Fe(III) and clay content (Fig. 3B). With repeated redox oscillations, crystallinity of aggregated Fe(III) may eventually reach a threshold at which root growth begins to be suppressed, marking the onset of SDF realized via negative root responses to crystalline Fe(III) (Fig. 2). However, since the processes that amplify Fe(III) aggregation form SDF by themselves, is clay-mediated SDF alone sufficient to form regular redox patterns? Or are both sets of SDFs required? If both, how do they interact during the pattern formation process?

**Figure 2.**
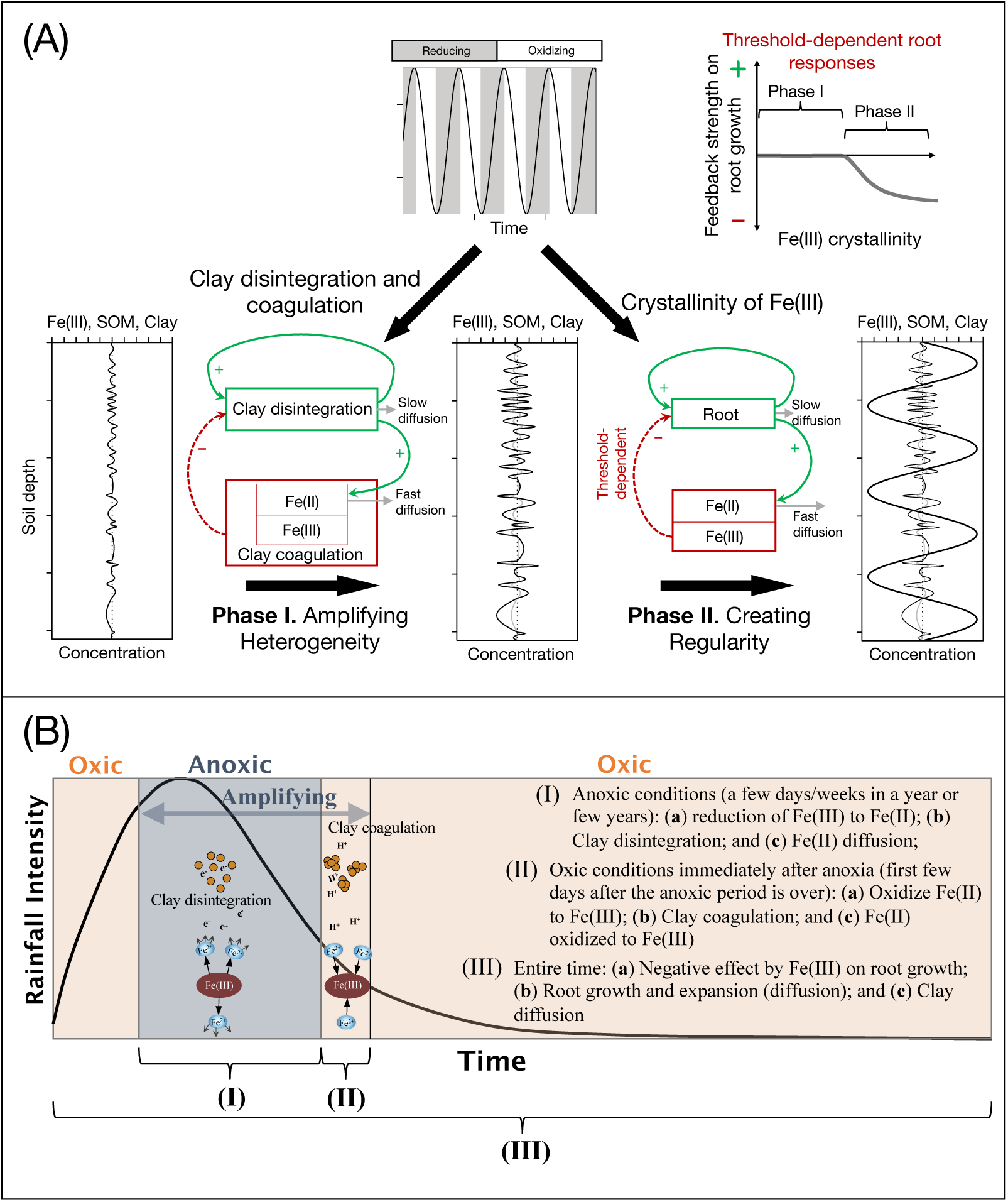
(A) Schematic of the two-phase mechanism underlying redox pattern formation in upland soils—the initial amplifying scale dependent feedbacks (SDF) phase and later threshold dependent SDF phase. Soil redox oscillations play an indispensable role in both phases. (B) Distinct timescales of key processes of SDF in soil redox pattern formation.

**Figure 3.**
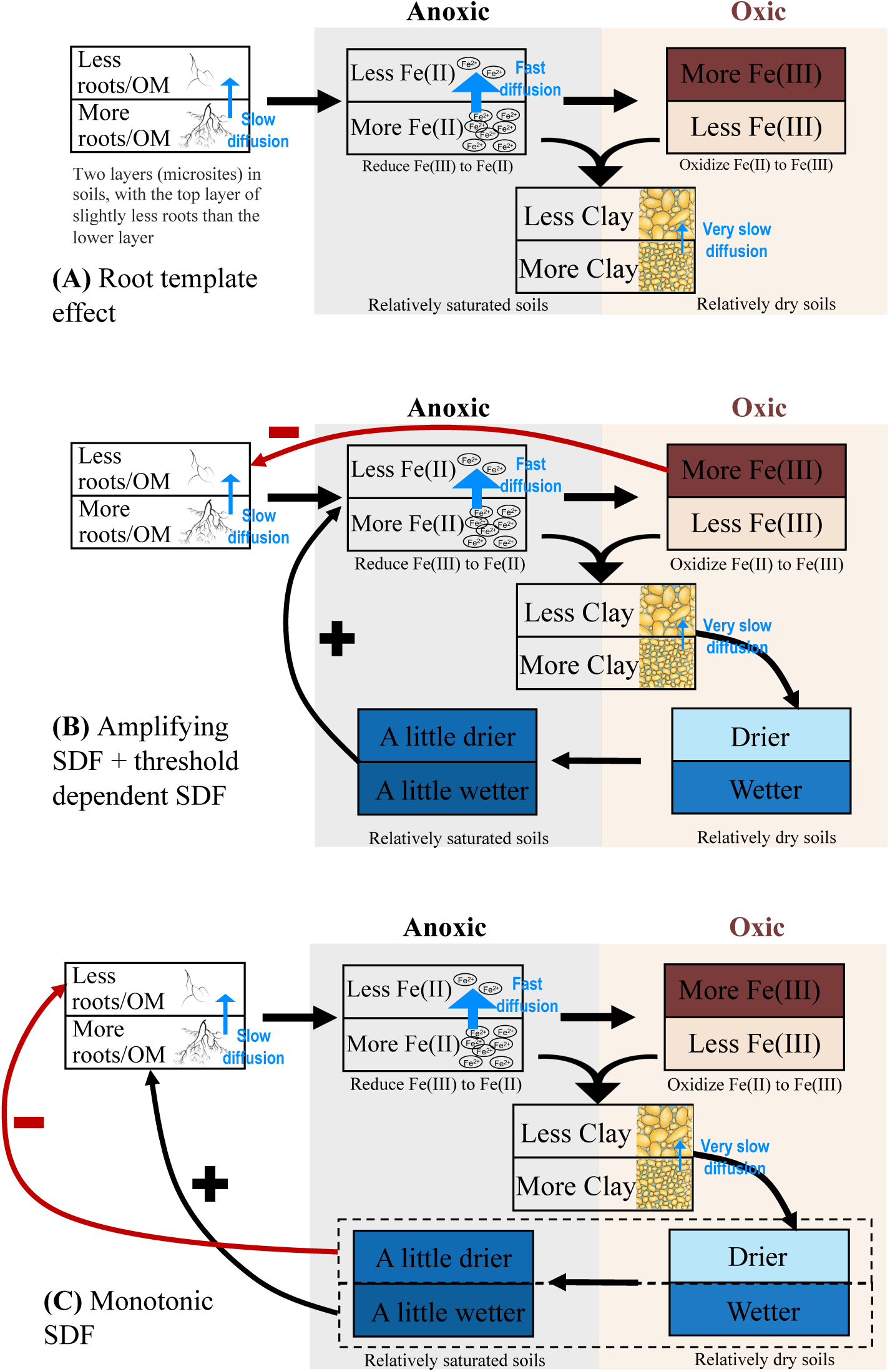
Three categories of hypothesized mechanisms giving rise to the regular redox patterns in upland soils. (A) Redox patterns formed as a reflection of the pre-existing root spatial structure. (B) Redox patterns formed as a coupling between amplifying SDF and threshold dependent SDF realized via a negative response of root growth to highly crystalline Fe(III). (C) Redox patterns formed by monotonic SDF mediated by root growth responses to soil water. Two stacked boxes represent two layers (microsites) of soils in the vertical direction, starting with the upper layer containing slightly less organic matter (OM), the lower layer containing slightly more OM, as the initial condition.

In this paper, we investigate mechanisms of redox pattern formation using a reactive transport model that couples chemical reactions, soil water dynamics, evolution of soil texture, and root growth. Our study site is at the Calhoun Experimental Forest, South Carolina, U.S.A. At this site, coarse-textured and porous A and E horizons overlie kandic Bt-horizons that are dominated by kaolinite and Fe and Al oxides. The area receives an annual rainfall of ∼ 1,270 mm. Rainfall periodically exceeds evapotranspiration (ET) and leads to low oxygen diffusivity, especially in B horizons, where a perched water table can emerge during winter and early spring months. When sufficient organic reductants are present, microbial decomposition consumes all of the oxygen and can switch to Fe(III) reduction as an alternative electron acceptor and mobilize Fe(II) via reductive dissolution (4, 40). At the Calhoun site, a pattern of alternating gray and orange bands, with band width of ∼ 1.6 cm (Fig. 1A and D), is located between the soil depths of 1.0 and 1.8 m (B horizons). Gray layers are low in Fe(III) but are rich in clay-sized minerals (kaolinite) and soil organic carbon (SOC) (although not higher than nearby sites without redox patterns) (Table S2). Gray layers harbor abundant fine roots and hyphal networks of mycorrhizal fungi. In contrast, orange layers are high in Fe(III) and low in clay-size particles and SOC, with no signs of root growth or ectomycorrhizal hyphae. Fe(III) crystallinity in orange layers is much higher than that in gray layers, with Fe_o_/Fe_d_ (ammonium-oxalate extractable Fe and dithionite-extractable Fe) ratio falling within the range for plinthite and ironstone concretions. A thorough description of physical and geochemical properties of redox patterns at our study site is provided elsewhere (4). In this study, we compare multiple, alternative mechanisms for pattern formation (Table S3). We will show that regular patterns in upland soils are generated by coupling two sets of SDFs—SDF based on clay dynamics and threshold-dependent SDF based on negative root responses to crystalline/plinthic Fe(III), with the former SDF amplifying Fe(III) aggregation and crystallinity to trigger the latter SDF. Temporal redox oscillations driven by environmental variability play an indispensable role in the formation of these banded redox patterns.

## 2. Results

### 2.1. Mechanisms of Pattern Formation

To sort among alternative mechanisms for patterns observed in the field, we examined three classes of mechanisms (in total five mechanisms/hypotheses; Table S3): (a) SDF with the negative feedback realized by *threshold-dependent* root responses to Fe(III), coupled with amplifying SDF mediated by clay dynamics (Video 1). This is our “baseline model.” Within this category of mechanism, we further investigated whether each set of SDF *alone* can give rise to the pattern (Video 2 and Video 3); (b) SDF with the negative feedback realized by *monotonic* root responses to water limitation (Video 4); and (c) template effect of pre-existing root structure determining the redox pattern (Video 5). Model simulated patterns are compared with empirically observed patterns to determine the most likely mechanism.

#### Pattern Set by Preexisting Root Structure

The most parsimonious mechanism to explain pattern formation is that the redox pattern is a manifestation of the pre-existing root spatial structure (Fig. 3A; Table S3). High SOM in the rhizosphere reduces Fe(III) to Fe(II), which diffuses away from rhizosphere, creating a gray zone surrounding the fine roots (Figs. 1G and 3A) and an orange zone surrounding the gray rhizosphere caused by enhanced deposit of Fe(III). If fine roots are already regularly spaced vertically, these processes can result in regularly spaced alternating gray–orange banded patterns. Using the evenly spaced root biomass as model initial condition, the steady-state redox pattern follows exactly root spatial distribution of the initial condition (Figs. S1 and S2). However, for redox patterns to follow the preexisting root distribution, it is required that roots do not expand into new regions (layers) of the soil (the diffusion coefficient of root biomass must be zero, *D_b_* = 0 in *Eq*. 8; Table S3) to allow a relatively stable template for redox patterns to form. Model simulations indicate that this mechanism would generate the banded pattern in ∼ 200 years.

#### Pattern Self-organized by Amplifying SDF alone

During redox oscillations, Fe(III) heterogeneity is amplified via clay-mediated SDF. Soil microsites of higher clay content retain higher moisture during dry-down and hence are more likely to be anoxic (Fig. 3B), leading to more Fe(III) reduction in clay-rich microsites. Increases in pH during Fe(III) reduction enhance disintegration of silt-sized kaolinite clay colloids, resulting in even higher clay content in high-clay microsites. Meanwhile, Fe(II) diffuses to nearby microsites of lower clay content that experience shorter periods of anoxia. In these low-clay microsites, Fe(II) oxidizes and precipitates as Fe oxyhydroxides and the lower pH associated with this process promotes clay coagulation and aggregation. Consequently, functional clay content declines in already low-clay microsites where Fe(III) continues to aggregate, while clay content increases in already high-clay microsites where Fe(III) continues to decline (*Eq*. 17). These processes amplify the pre-existing degree of heterogeneity of clay and Fe in soils.

While serving as an amplifier, we found that clay mediated SDF can also give rise to regular patterning (Fig. S1; Table S3); however, under the rainfall regime at our study site, formation of regular patterning by this mechanism requires a very slow rate of bulk clay diffusion. Clay disintegration (short-range positive feedback in SDF) and coagulation (long-range negative feedback) in this model only occur under anoxic conditions and only for a few days after soil anoxia begins. This is a very brief period over the course of several years. In contrast, clay diffusion occurs year around (Fig. 2B). A large clay diffusion coefficient (*D_c_*) would smooth out clay heterogeneity, preventing pattern formation. Under the climatic regime of our study site, only when *D_c_* is < 2.5×10^-16^ m^2^ s^-1^, can regular patterns arise; however, the emergent banding width is only ∼ 1/3 of the width observed in the field. While it is possible to obtain banding widths comparable to the observed width by increasing *D_c_*, for patterns to still form under a larger *D_c_*, a much wetter climate is required, allowing for more frequent or longer anoxic conditions. Longer or more frequent anoxic conditions increase *mean annual* rates of Fe(III) reduction and oxidation; that is, greater clay disintegration and coagulation are required to counteract the smoothing caused by year-round diffusion.

Under the climatic condition of our study site, the formation of narrow banding requires ∼ 7,000 years, an order of magnitude longer than the estimated time required by other mechanisms (Table S3). Long pattern formation time is caused by the amplifying SDF being weak and slow (Fig. S3). The nature of the amplifying effect is differential soil moisture content (*Se* in *Eq*. 4) in gray and orange layers. This results in distinct clay dynamics—clay disintegration dominating in gray layers and clay coagulation in orange layers. Larger differences in soil moisture content between orange and grey layers create stronger amplifying effects. However, the key process underlying the amplifying SDF, Fe(III) reduction, requires anoxic conditions, which are not common in upland soils (Fig. S4). More importantly, when the anoxic requirement is finally met in (nearly) saturated soils (DO < 0.02 mol m^-3^), values of soil moisture in orange and gray layers are similar (Fig. S3); i.e., all layers are close to saturation. These conditions dictate that the amplifying effects that enhance Fe(III) aggregation and crystallinity are weak.

Furthermore, with amplifying SDF alone, simulated OM content is slightly higher in the orange layers than in the gray layers (Fig. S1-C5), contrary to the observed (field) pattern showing significantly lower OM in orange layers. Under oxic conditions, a similar amount of OM is oxidized by DO in both orange and gray layers. Under anoxic conditions, slightly more OM is oxidized by Fe(III) in gray layers where normalized soil water content (*Se* in *Eq*. 4) is higher. Without a negative effect on root growth by resource limitation or inhibitors, as hypothesized in other mechanisms, a similar amount of OM is produced in both orange and gray layers. As a result, lower OM consumption in orange layers generates slightly higher OM in orange layers, a pattern inconsistent with that observed in the field (Fig. S1; Table S3). Because multiple aspects of model results (band width and OM pattern) contradict field-observed patterns, amplifying SDF alone are unlikely to be the mechanism responsible for formation of the observed banded redox patterns.

#### Pattern Self-organized by Threshold-dependent SDF Coupled with Amplifying SDF

Amplifying SDF described above increase heterogeneity of Fe(III). Over repeated redox oscillations, Fe(III) aggregation and crystallinity intensify and eventually reach a threshold when an array of changes occurs and the negative effects on root growth begin. This triggers the second set of Turing instability (Fig. 2A). Soils with more crystalline Fe(III) or plinthic-like horizons often have diminished nutrient and water retention properties (31, 32, 41) and these can limit root growth. Additionally, highly crystalline Fe(III)— plinthite, often observed in soil redoximorphic features, is hard and brittle, forming a barrier to root penetration (27, 33). When lower crystallinity Fe(III) dominates the Fe pool, roots usually can effectively penetrate soils by various morphological and chemical adaptations (42, 43). At our study site, goethite is present in both gray and orange layers; however, its crystallinity is significantly higher in orange layers (44), and thus fine roots grow exclusively in gray layers (4). Such a root growth pattern is found in other places with similar redox banding (5). Radiocarbon dating at our study site shows that SOM in gray layers is significantly younger than that in orange layers, indicating lack of recent root growth in orange layers (4). Based on these lines of evidence, we used a threshold-dependent function (*Eqs.* 10 & 11; Table S3) to describe the effect of Fe(III) on root growth. Fe(III) is a surrogate for an array of changes occurring in soils as Fe(III) aggregation and crystallinity both increase over repeated redox oscillations.

Slow diffusion (expansion) of roots (an activator in the Turing instability framework) and rapid diffusion of Fe(II) (an inhibitor of root expansion in oxidized form) give rise to the regular pattern. Once triggered, the threshold dependent SDF coarsens the narrow banding formed by the amplifying SDF, generating banding widths similar to observed values. The model that couples the two sets of SDFs reproduces *all* aspects of spatial patterns in the field (Figs. 4 and S1; Tables S2 and S3), and ∼ 900 years is required for patterns to form. This is the most likely explanation for redox patterning observed at our study site (Table S3). When the amplifying SDF is turned off in the model simulation, leaving threshold dependent SDF to operate alone, no banded patterns arise (Fig. S1; Table S3).

**Figure 4.**
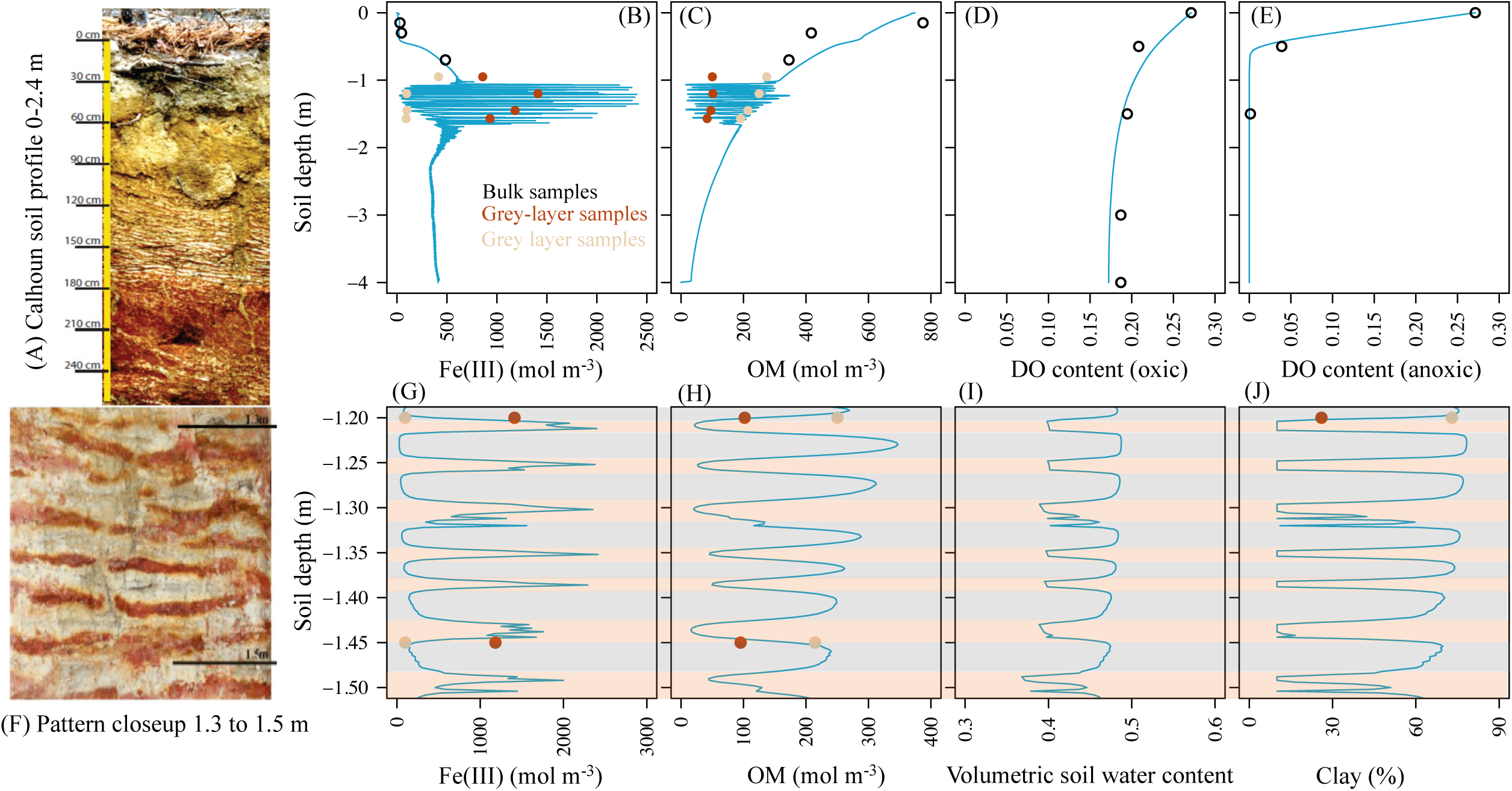
Comparison between modeled and observed soil redox patterns. (A) Photo shows the regular iron banding at the study site at the Calhoun Experimental Forest, with the close-up view of the redox pattern between 1.3 and 1.5 m soil depth (F). (B)-(E) blue lines show model simulated Fe(III) concentration (B), OM concentration (C), dissolved oxygen (DO) content under oxic condition (D) and DO content under anoxic condition (E). Points in each plot denotes measurements from the study site, with dark brown representing samples from orange layers, and light-yellow representing samples from gray layers. Open black circles represent bulk samples. (G) – (J) are close-up view between 1.2 and 1.5 m soil depth of the model simulated patterns of Fe(III) concentration (G), OM concentration (H), soil volumetric water content (I), and clay content (J). (G)-(J) show that layers of high Fe(III) show low OM, low soil water, and low clay content, while layers of low Fe(III) show high OM, high soil water, and high clay content. The DO profile in oxic condition (D) and volumetric water content (I) is model result after 30 days of exposure to aerobic conditions.

#### Pattern Self-organized by Monotonic SDF alone

We further tested the hypothesis of a *monotonic* relationship between root growth and resource level, specifically of soil water. That is, higher water content stimulates root growth while lower water content restricts growth (Table S3). Unlike the mechanism of threshold-dependent inhibitory effects of Fe(III) on roots, the model variant with this mechanism (monotonic responses of root growth to soil moisture) can reproduce regular patterns *without* amplifying SDF (Fig. S1; Table S3). During anoxic episodes, in microsites of high OM, a large amount of Fe(III) is reduced, with marked clay disintegration, increasing clay content and soil water holding capacity, which in turn stimulates root growth (Fig. 3C). More root growth leads to more SOM accumulation, accompanied by Fe(III) depletion during subsequent anoxic periods. Meanwhile, Fe(II) from these microsites diffuses and is oxidized to Fe(III) in nearby orange layers during oxic episodes. Oxidative precipitation facilitates clay coagulation, reducing clay content when Fe(III) precipitates. In microsites of low clay, soil water content is relatively low and can limit root growth (Fig. 3C). The model with this mechanism can reproduce Fe(III) and clay patterns; however, the model predicts much higher SOM in gray layers than is observed in the field (Fig. S1), resulting from positive root responses to the relatively high-water content in clay-enriched gray layers. At our study site, SOM content in gray layers is not higher than it is in those nearby sites without pattern formation (indicative of background level; Fig. S1). This suggests a minimal to non-existent stimulating effect on root growth in gray layers. Pattern formation occurs in ∼ 900 years when this mechanism applies.

### 2.2. Conditions for Pattern Formation

One of the most important conditions for pattern formation is redox fluctuations, which is largely determined by climate and soil conditions (texture and labile carbon availability). Regular patterns are more likely to form in relatively dry conditions and in soils with an intermediate clay content (Fig. 5). Under the parameterization of our model, regular patterns are likely to emerge when precipitation is < 3,500 mm yr^-1^ and clay content is between 40% and 80%. Both wet conditions and high clay content can enhance Fe leaching, reducing the likelihood of patterning. While soils of high clay content increase leaching, soils of low clay content feature high hydraulic conductivity and low water retention capacity (Figs. S5 and S6), often resulting in well-oxygenated soils, limiting the soil anoxia needed for pattern formation. Furthermore, when annual precipitation increases, the pattern-forming range of clay content narrows and lowers (Fig. 5), suggesting a compensatory effect of a higher clay content for a drier climate. Such a compensatory pattern is also observed empirically in data from sites in different parts of the world (Fig. 5).

**Figure 5.**
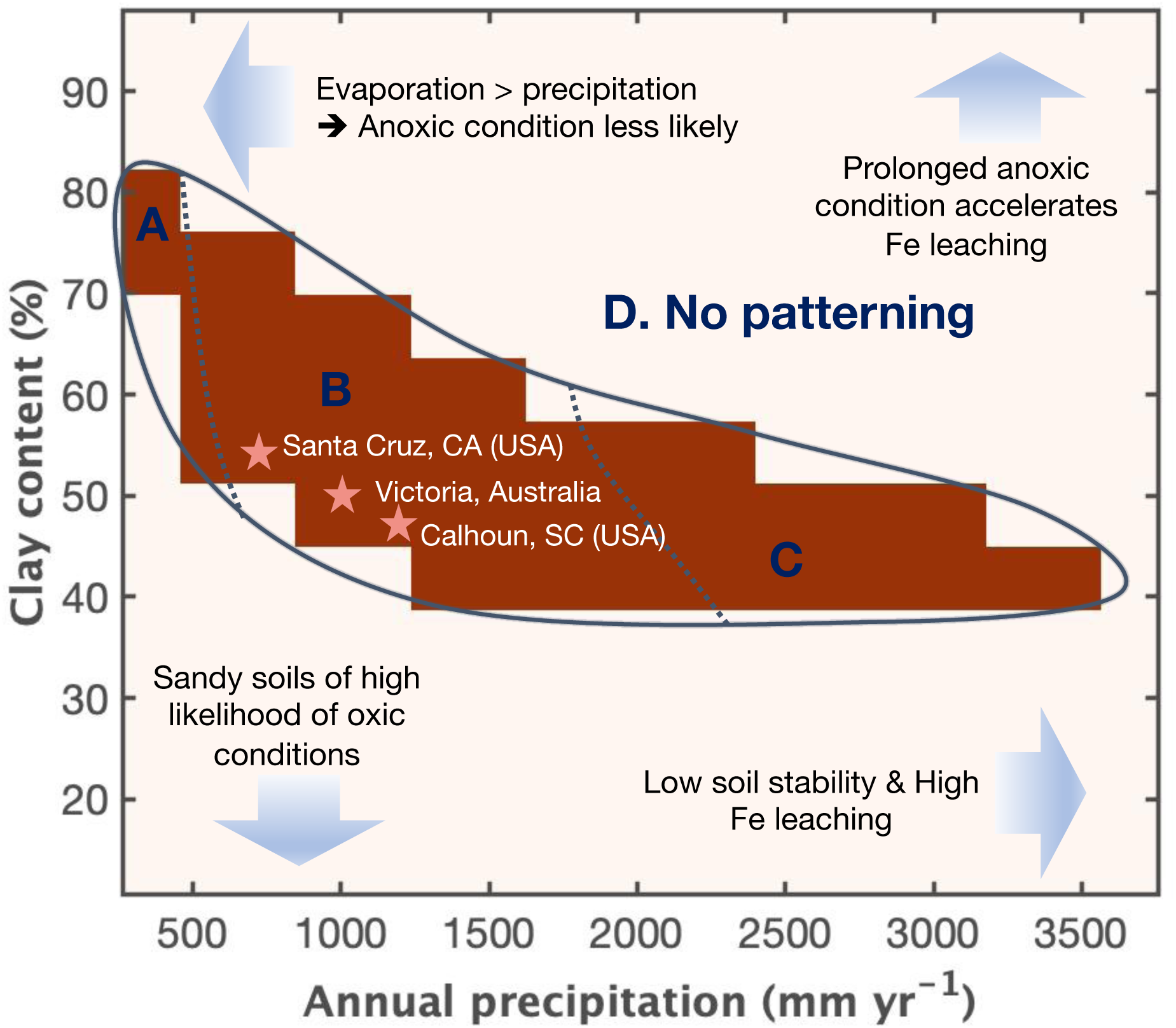
The domain of soil texture and annual precipitation where regular redox patterns are likely to emerge in upland soils. The solid line polygon delineates the domain predicted by the model where regular redox patterning will form. Zone D depicts the domain without patterning. Within the solid-line polygon, we expect pattern formation is most likely to occur in Zone B. The model manipulates different levels of annual precipitations but assumes same evaporation rate. If temperature increases in more arid zones, regular patterns are unlikely to emerge in Zone A. The model does not consider soil stability; however, if soils become less stable in wetter regions, it is unlikely to form regular redox patterns (Zone C). The stars indicate the condition of soil and precipitation at three sites where regular redox patterns have been reported (Table S3).

The upper and lower boundaries of the pattern formation zone are determined by the vertical distributions of clay, Fe(III), and OM in the soil. Regular patterning starts at 40 – 50 cm below the location where clay content begins to rapidly increase (Figs. 4 and S7D). This is the soil depth where anoxic conditions are more likely due to subsoil water perching (Figs. 4 and S4) and high Fe(III) (Fig. S7C). Soils above this zone are sandy and almost always oxic (Figs. S7), except in the early spring when high OM decomposition limits O_2_ diffusion (45). At deeper soil depths (> 2.5 m), OM becomes too low for the Fe(III) reduction required for pattern formation. Sensitivity of the Fe(III) reduction rate to soil water content also affects the upper and lower boundaries of the pattern formation zone (Fig. S3). When sensitivity is low, only the region with relatively high Fe(III) can reach the threshold of negative feedbacks, resulting in narrow pattern formation zones (Fig. S8). In contrast, if the rate of Fe(III) reduction is highly sensitive to soil water content, the pattern formation zone expands. The width of pattern elements (banding) is primarily controlled by the two diffusion coefficients, i.e., the diffusion coefficient for Fe(II) (*D_10_* in *Eq*. 7) and for root biomass (*D_b_* in *Eq*. 8) (Fig. 6). Banding width increases with diffusion. Our models suggest that for a pattern to emerge, *D_b_* must be at least three orders of magnitude smaller than *D_10_*. Further, the clay diffusion coefficient *D_c_* (*Eq*. 17) must be even smaller than *D_b_*.

**Figure 6.**
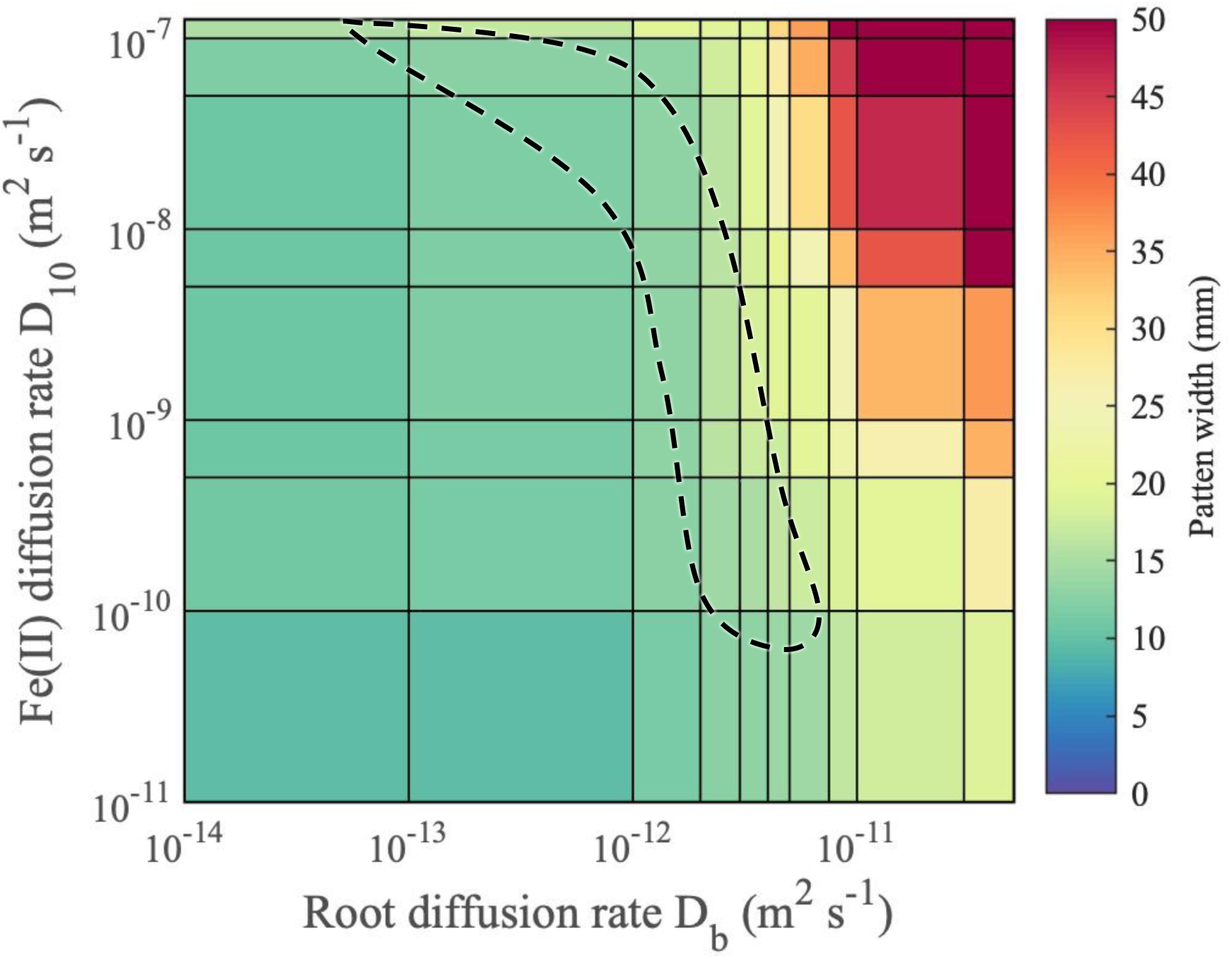
The effect of diffusion coefficients of root biomass (*D_b_*) and diffusion coefficient of Fe^2+^ in soil water (*D_10_*) on the width of the regular redox patterns in soil. The dash-line polygon delineates the area where pattern width (average width of an orange layer or a gray layer, ∼ 1.6 cm) is representative of that at our study site.

The time required for patterns to form is highly dependent on climate. In our study system, the anoxic period occurs in winter and spring, when evaporation rate is relatively low (Fig. 7) (45). Our baseline model (the model coupling two sets of SDFs; Table S3) estimates that ∼ 900 years is required for regular patterns to form. Given that the annual precipitation regime imposed on the model is approximately at the 4-year recurrence interval, the actual pattern formation time is near ∼ 3,600 years. A much longer time is required for patterns to form in drier areas because the likelihood of wet conditions that induce soil anoxia is much lower. In fact, we found that with a decline in annual precipitation, required pattern formation time increases *exponentially* (Fig. S9). The time required for patterns to form is further affected by the rate of Fe(III) reduction in an approximately linear fashion (Fig. S9).

**Figure 7.**
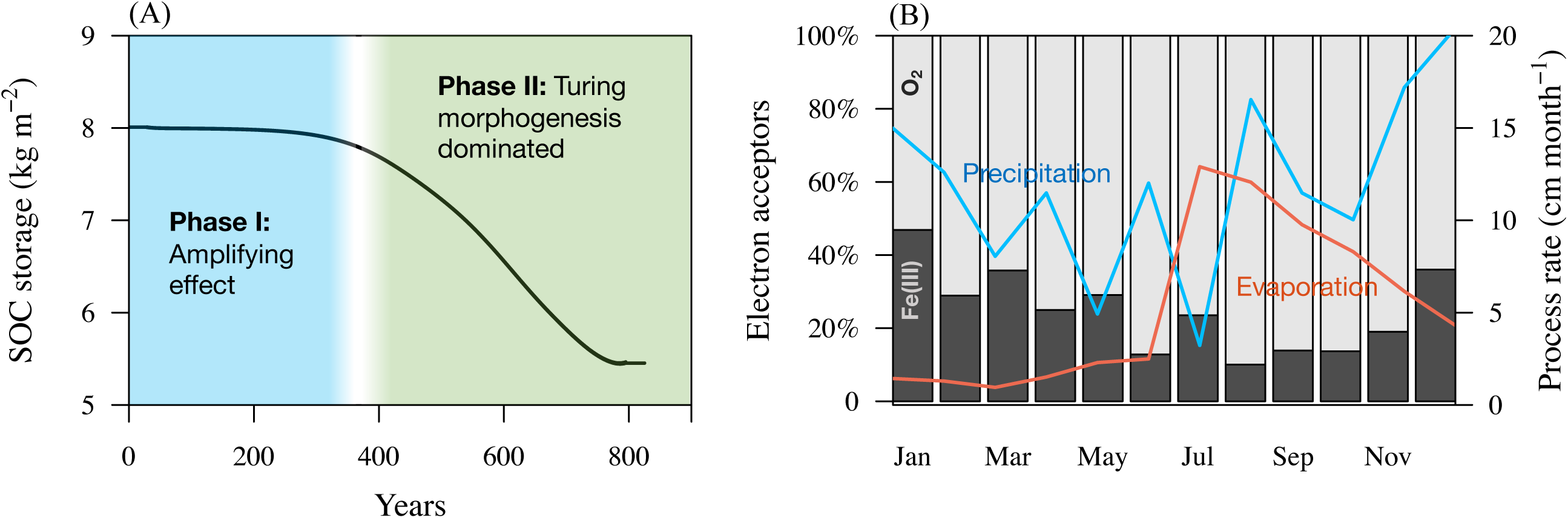
(A) A reduction of ∼30% in the soil organic carbon (SOC) storage capacity in the pattern formation zone (between 1.0-1.8 m of soil depths) over the 900-year pattern forming period. (B) Percent of SOC oxidized by O_2_ *vs.* by Fe(III) by month in the model parameterized by the environmental conditions of our study site in a representative wet year at the Calhoun Experimental Forest (South Carolina, U.S.). Data of monthly rainfall and evaporation rate (cm per month) are aggregated from the daily rate monitored at the study site between July 1, 2018 to June 30, 2019.

### 2.3. Consequences of Pattern Formation for Soil Carbon

Since pattern formation involves root dynamics, redox patterning might affect soil organic carbon (SOC), both in terms of its storage capacity and flux. At our study site, orange layers are ∼ 2/3 the width of the gray layers, and SOC content in the orange layers is ∼ 40% of that in gray layers. If we use SOC in gray layers—which is similar to SOC content in nearby sites without regular patterns—as the background level for the region, formation of orange layers results in a 24% reduction in SOC storage within the pattern formation zone (1.0-1.8 m). Our model reproduced this amount of carbon storage reduction by pattern formation (Fig. 7). The initial amplifying SDF has little effect on soil carbon storage, but when crystalline Fe(III) starts to exert a negative effect on root growth, carbon storage is reduced. At our study site, SOC in the pattern-forming zone accounts for ∼ 20% of total SOC in the top 4 m of soils (model domain). This means that pattern formation reduced the carbon storage capacity of the top 4 m of soils by 4.8%. For the total SOM oxidized each year, 10 – 50% is oxidized by Fe(III), and the remainder by O_2_ (Fig. 7B). Fe(III) serves as an important electron accepter between December and May, the period of low evaporation, although precipitation is not high during that time of a year (Fig. 7A). As a result of the spatial segregation of OM and Fe(III) in the gray and orange layers created by pattern formation, a potential capacity of ∼ 0.86 t C ha^-1^ mineralization by Fe(III) is lost (estimated from (46)).

## 3. Discussion

### Mechanism of Pattern Formation

Turing instability forms regular patterns under conditions of environmental fluctuation. However, some inhibitors may need to reach a certain level or state (e.g., a plinthite-like form of Fe(III)) to achieve Turing instability. This makes the inhibitory feedback threshold-dependent. Threshold-dependent responses are common in nature (47–49); however, their effects on Turing morphogenesis have seldom been investigated. The most plausible mechanism of redox patterning in upland soils is the coupling of two sets of SDFs—clay-mediated SDF amplifies Fe(III) aggregation and crystallinity to the state that suppresses root growth. This then triggers the second set of SDF. Plants develop different approaches to improve their root penetration in soils (42, 43). It is not until Fe(III) increases in crystallinity and forms plinthite-like aggregates, that these approaches would become ineffective, triggering negative growth responses. In the orange layers where SOM is low, Fe(II) can promote the structural transformation of weakly crystalline Fe(III) into more crystalline and thermodynamically stable phases of Fe(III) (50, 51), which can then serve as a template for greater precipitation of crystalline-Fe(III) (52). Experiments using soils from our study site with high native Fe(III) crystallinity and depleted SOM produced goethite with exceptionally high crystallinity (52). In contrast, in gray layers, enriched SOM inhibited the transformation of weakly crystalline Fe(III) into a more crystalline form (50). At our study site, goethite in SOM-enriched gray layers has a much lower crystallinity than it does in the SOM-depleted orange layers (44). Coupled SDFs are likely generalizable to many self-organized systems where SDF are threshold dependent, a functional form common in nature (47–49).

The nature of processes that trigger negative growth responses is SDF as well. These can give rise to regular patterns by themselves. However, amplifying SDF is weak (Fig. S3), resulting in an order of magnitude longer time for patterns to form by this mechanism alone (Table S3). Importantly, because the amplifying SDF is relatively weak, they do not strongly interfere with the second set of SDF, which set the characteristic band width. Otherwise, the pattern will not be regularly spaced, with different band widths set by different sets of SDFs. SDF based on *monotonic* root responses to soil water content can give rise to patterning even without the amplifying phase. As monotonic growth responses to soil water content assume a positive root response to high water content, the simulated SOM in gray layers becomes much higher than field observations suggest (Fig. S1-E4). Previous studies have shown that positive feedbacks in SDF are not a necessary condition for pattern formation (53). Therefore, long-range negative feedback alone would produce patterns that match field observation, if the negative feedback can be effective at the initial condition, i.e., the initial soil moisture is low enough to suppress root growth. However, soil moisture at our study site is high overall, averaging ∼ 38% to 43% at 0.5 m depth year around (Fig. S4) and root growth is not likely to be suppressed at such a high moisture level (54). In that case, for the negative feedback alone to create the pattern, additional processes that can amplify the spatial heterogeneity of soil moisture are required so that local moisture can be sufficiently low to suppress root growth. This moisture mediated monotonic SDF might contribute to redox pattern formation in much drier soils. Monotonic plant growth responses to water availability has explained regular vegetation patterns in drylands around the world (53, 55).

The root template hypothesis—that redox patterns are determined by the pre-existing root structure—is also unlikely to explain observed patterns. If soil redox patterns are a mere reflection of preexisting root structure, this mechanism would require (I) a regularly spaced lateral root system; and (II) the same root spatial structure persisting for a long enough time to allow pattern formation. Neither condition is likely. Observations of root distributions do not show a regular structure that mirrors the observed redox patterns (Fig. S10). Even if such regular structure exists, fine roots are deciduous (56) and are highly dynamic in time so as to allow plants to rapidly adjust root structure to compete for limiting resources (57). Furthermore, while the same loblolly pine species dominates most of the study area, regular patterns are very patchy, indicating other drivers than plants are important. Nevertheless, many irregular redoximorphic features in soils follow exactly the root structure, showing an iron-depleted gray rhizosphere in upland soils (Fig. 1G) and an iron-concentrated orange rhizosphere in hydric soils (“mottles”) (Fig. 1F) (4, 5, 58).

Beside SDF, phase separation can also drive spatial self-organization (9, 19). When a binary mixture is subject to periodic forcing, e.g., freeze-thaw cycles in cold regions, particles autogenically separate and form distinct spatial patterns (19, 59). The concentration-dependent movement feedback—switching from dispersion to aggregation as local concentration increases—is central to the phase separation principle and subsequent pattern generation. For example, mussels move at a high speed to form clusters when they are at low density and once incorporated in clusters, they decrease their speed of movement. This density-dependent movement feedback generates the diverse self-organized patterns of mussel beds (9, 60). In our case, although redox cycles lead to local aggregation of clay, this does not occur through concentration-dependent clay movement. Instead, clay aggregation occurs by disintegration in microsites locally, without translocation (37).

### Why Regular Redox Patterning might be Rare?

While redoximorphic features are common in soils, rhythmic redox banding is rare. This likely has to do with the intricate coordination of climatic, soil, and biological conditions required for redox pattern formation. Upland soils are often well-aerated. In subtropical forests, such as Calhoun in the Southeastern U.S. Piedmont, the marked seasonality of rainfall, transpiration, and temperature restricts anoxic conditions to only short periods within wet years (45, 46) (Figs. 7 and S4). Soil moisture can be further modified by local factors, e.g., land management and topography. As such, even on the same landscape, regular patterns can be highly patchy. For instance, no redoximorphic features appear in a soil pit only 400 m from our study site. Another pit ∼ 4,000 m away shows sparse, irregular patterns (Fig. 1). While anoxic conditions are more likely in wetter environments, soil stability declines in wetter environments (61). As pattern formation likely requires thousands of years, the likelihood of pattern formation is reduced when soil stability is low. In addition, prolonged anoxic periods in wet environments can accelerate Fe leaching, especially in soils with abundant organic acids (62), reducing the likelihood of pattern formation (Fig. 5). Eventually inundation can be detrimental to upland plants (63). Plants that adapt to extended inundation, e.g., wetland plants, can transport O_2_ to their roots. A proportion of O_2_ diffuses to the rhizosphere, forming iron plague on the root surface by oxidative precipitation (64, 65) (Fig. 1F). This is in contrast to the iron depleted gray rhizosphere of upland plants created by reductive dissolution (5) (Fig. 1G). If the wetland soil contains a reservoir of Fe(II), regular banding of Fe(III) precipitate might emerge via the same mechanism that forms Liesegang bands in rocks. A classic Liesegang mechanism involves a dissolved reactant diffusing from a system’s boundary into a reservoir of another reactant. Subsequent precipitate banding forms in the wake of a propagating reaction front (66). Roots however represent distributed O_2_ sources, and disrupt the operation of self-organization. We speculate that redoximorphic features in wetlands are in fact likely set by the root template.

Formation of rhythmic redox patterns requires the spatial alignment of vertical distributions of root growth, clay, and Fe(III). Pattern formation occurs in zones of high clay and Fe(III) (Fig. S7). Clay accumulations in soils often occur between 0.3 and 3 m depths (67), a range determined by parent material, soil developmental stage, hydrologic regime, land use, etc. (68–70). Similar to clay-enriched horizons, Fe profiles are also sensitive to many factors, e.g., groundwater table, topography-driven surface flow (71). Furthermore, pattern formation requires fine root proliferation at relatively deep soil layers, overlapping vertically with the high-clay, high-Fe subsoils. At our study site, fine roots of the dominant species, loblolly pine, are present even at 4-m depth (68). B horizons, characterized by high clay and Fe, are the most likely zone for pattern formation. Reported regular redox banding from different regions of the world all occur in B horizons (4–6) (Table S1).

### Temporal Variability in Pattern Formation

Environmental variability is an essential feature of many ecosystems around the world; however, the general role of environmental variability on ecosystem pattern formation is so far been under-studied. In cold environments, freeze-thaw cycles create feedbacks essential for the mechanism of phase separation, producing self-organized sorted circles (17, 19, 59). For spatial self-organization by Turing instability, redox patterns in the upland soils studied here exemplify the significance of temporal dynamics. Environmental variability produces diverse conditions that allow various processes required for pattern formation to occur at the same site, although in different temporal windows orchestrating over the timescale relevant for pattern formation. Temporal oscillation itself constitutes a unique condition that allows certain processes to occur. For example, repeated redox changes induce clay disintegration/coagulation and enhance phases of Fe(III) crystallinity (26, 37, 51). Redox variability plays an indispensable role in both sets of SDFs that give rise to soil redox patterns (Fig. 2).

Environmental variability further affects mean process rates over timescales relevant to pattern formation. Turing morphogenesis requires the inhibitor to diffuse faster than the activator (Fig. 2A), allowing the resultant long-distance negative feedback to suppress the expansion of the activator (11, 53). However, what is important is their *mean* rates over timescales relevant to pattern formation. Pattern formation in upland soils couples chemical, physical, and biological processes at distinct timescales. Some processes operate for only a brief period over a span of a few years, while others operate continuously. Temporal intermittency of SDF processes, driven by environmental variability, significantly alter mean annual process rates, thereby modifying the likelihood of pattern formation. In our study, roots act as the activator and expand throughout the year. However, Fe(III) reductive dissolution, Fe(II) transport, and oxidative precipitation occur intermittently, perhaps for only a few days or weeks over several years in upland soils (Fig. 2B); that is, the diffusion of the inhibitor, Fe(II), operates for a much shorter period than the activator (roots). Under the climatic regime of our system, for regular patterns to form, we find that root diffusion must be at least three orders of magnitude slower than Fe(II) diffusion (Fig. 6). When the environment is drier, an even smaller rate of root diffusion is requisite. Fine root extension of the dominant species *Pinus taeda* at our study site is slow compared to many other common subtropical trees (72). The root diffusion coefficient of *P. taeda* is ∼ 1.8×10^-11^ m^2^ s^-1^, estimated by the method in (73) using empirical *P. taeda* fine root extension rates (74). This is surprisingly close to our model-calibrated value (Table S4). Similarly, in the amplifying SDF, clay disintegration and coagulation, and the transport of inhibitor (Fe(II)) occur for only brief periods over one or several years; however, clay diffusion runs continuously. As diffusion smooths out heterogeneity, clay diffusion rate must be very low for effective amplification. The calibrated clay diffusion coefficient, comparable to experimental measurements (75), is several orders of magnitude lower than that of root diffusion (Table S4).

This study illustrates the critical role played by environmental variability in forming soil redox patterns. Environmental variability prevails in ecosystems worldwide, directly controlling the operation of a wide range of chemical and physical processes and the timing and rates of many biological processes (e.g., phenology, life cycle). Given the significant role of temporal dynamics in pattern formation, projected changes in climatic variability might substantially alter many self-organized systems, including important ecological consequences associated with the striking patterns they display.

## METHODS

### Modeling Redox Reactions in Upland Soils

We constructed a reactive transport model coupling soil water dynamics and iron redox reactions to investigate the mechanism of redox pattern formation. The spatial domain of the model extends from the soil-air boundary to 4 m soil depth. Model spatial resolution is 2 mm with a time step of 1 hour, to capture the fine-scale spatial pattern and rapid chemical processes respectively. The part of the model describing soil redox reactions includes six state variables, oxidized iron Fe(III), reduced iron Fe(II), root biomass *B*, organic matter *OM*, clay, and oxygen concentrations, including dissolved oxygen *O_2L_* and gaseous oxygen *O_2G_*. Model parameters, their definitions, dimensions, and choice of values are provided in Table S4.

Fe(III) reductive dissolution and oxidative precipitation can be described by the following reactions:

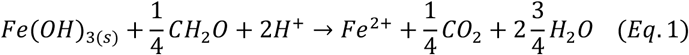

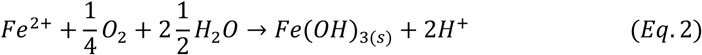

Therefore, the change of Fe(III) concentration (mol m^-3^) is expressed as follows:

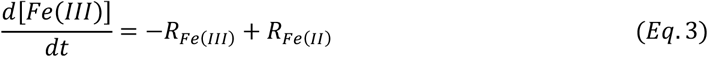

where *R_Fe(III)_* (mol m^-3^ d^-1^) is the rate of Fe(III) reductive dissolution (*Eq*. 1) and *R_Fe(II)_* (mol m^-3^ d^-1^) is the reaction rate of Fe(II) oxidative precipitation (*Eq.* 2). We assume the rate of Fe(III) reductive dissolution is controlled by concentrations of *O_2L_*, *OM*, Fe(III) and soil water content (76–78). When soil *O_2L_* is high, reductive dissolution is negligible (79). Reductive dissolution of Fe(III) becomes significant when *O_2L_* reaches a very low level (0.02 mol m^-3^). Fe reduction is further positively affected by soil water content, which determines the amount of low-oxygen microsites available in soil matrix (38) (Fig. S11). Rate of Fe(III) reductive dissolution is described as:

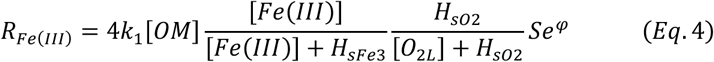

where [*OM*] is the concentration of SOC (mol m^-3^), [*O_2L_*] is the concentration of *O_2L_* (mol m^-3^), [Fe(III)] is the concentration of Fe(III) (mol m^-3^), and *k*_1_ is the constant of the first-order organic matter reductive rate (d^-1^). *H*_*sFe*3_ (mol m^-3^) is the half saturation constant regulating the effect of Fe(III) on the reduction rate, and *H_so2_* (mol m^-3^) is the half-saturation constant regulating the effect of *O_2L_*. *Se* is normalized volumetric water content, which varies in space and time. Dynamics of *Se* is described in *Eq*. 19 in Section “*Soil water dynamics*” below. *φ* is a constant describing the sensitivity of the rate of Fe(III) reduction to *Se*.

The rate of Fe(II) oxidative precipitation *R_Fe(II)_* (mol m^-3^ d^-1^) can be described as:

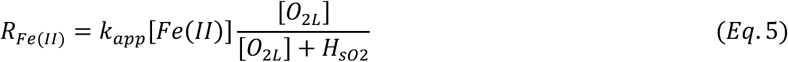

where [Fe(II)] is the concentration of Fe(II) (mol m^-3^); *k_app_* is the constant of Fe(II) oxidation rate (d^-1^). Fe(II) is affected by chemical reactions, advection, and diffusion:

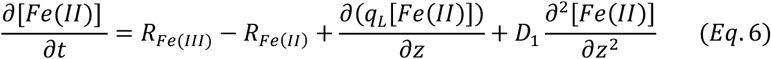

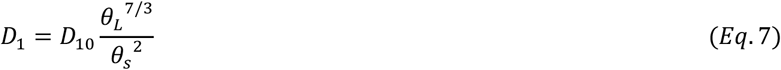

where *q_L_* is soil water flux (m d^-1^) and *D_1_* is the diffusion coefficient (m^2^ d^-1^) of Fe(II) in soil, which varies with soil water content as described in *Eq*. 7 (80). *D_10_* is the molecular diffusivity of Fe(II). *θ_L_* is soil water content. Note that *θ_G_* changes in space and time, whose dynamics is modeled in the Section “*Soil Water Dynamics*” below.

Change of root biomass density [*B*] (g m^-3^) is affected by (1) root growth, *G_B_*, (2) root respiration, *R_B_*, and (3) root decay, *M_B_*. In addition to root growth, which describes biomass increases in a given location, roots can also expand to neighboring locations. Root expansion in space is commonly approximated by a diffusion term (55, 73):

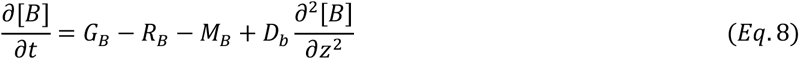

where *D_b_* is the diffusion coefficient (m^2^ d^-1^) of root biomass. Root respiration *R_B_* (g m^-3^ d^-1^) is influenced by root biomass density and soil oxygen concentration:

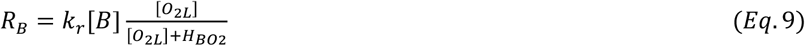

where *k_r_* is root respiration rate constant (d^-1^). *H_BO2_* is half-saturation constant to regulate the effect of *O_2L_* on root respiration (mol m^-3^). Root growth could be influenced by an array of factors, such as soil water content, nutrients and oxygen availability, and mechanical impedance (81, 82). We describe root growth *G_B_* (g m^-3^ d^-1^) as follows:

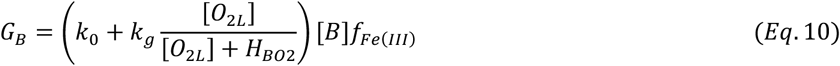

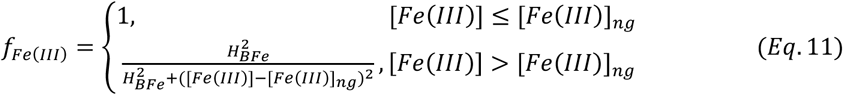

where *k_0_* (d^-1^) is the root growth rate constant, and *K_g_* (d^-1^) is that part of root growth affected by oxygen. *f_Fe(III)_* is a scaler, describing the effect of Fe(III) on root growth, ranging between 0 and 1. We use Fe(III) as a surrogate to describe integrated effects of multiple environmental factors associated with Fe(III) aggregation on root growth. While high Fe(III) concentrations do not lead to increases in crystallinity *per se*, under repeated redox oscillations over time, increases in Fe(III) aggregation and in the crystallinity of Fe(III) usually coincide (26, 83). Aggregated Fe(III) precipitates can lead to coarse soil texture, low soil water content, and low nutrient availability, and highly crystalline Fe(III) aggregates (e.g., plinthitic material) create high mechanical impedance for root growth (4, 32, 33, 84). [*Fe(III)*]*_ng_* denotes the threshold Fe(III) concentration (mol m^-3^), above which Fe(III) will negatively affect root growth. *H_BFe_* is the half-saturation constant regulating the effect of Fe(III) on root growth (mol m^-3^). Root decay, *M_B_* (g m^-3^ d^-1^), is assumed to be a second-order function of biomass (84):

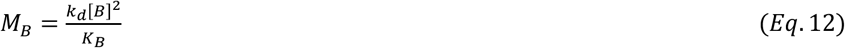

where *k_d_* is root decay constant (d^-1^), and *K_B_* (g m^-3^) is the carrying capacity of root biomass density, which decreases exponentially with soil depth (Table S4; Fig. S10). We assume that root decay is the primary source of SOM and SOM is oxidized by O_2_ or Fe(III). We note that the root biomass, *B*, is a lumped variable including roots, mycorrhizal fungal symbionts, and microbes, all contributing to the SOM pool (85). The change of SOM is described as:

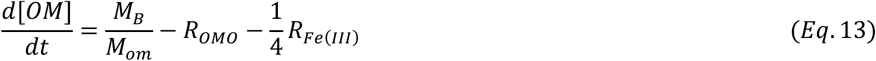

where *M_om_* is molar mass of SOM, assuming 30 g mol^-1^ for the generic formula of CH_2_O. Rate of SOM oxidation by O_2_, *R_OMO_* (mol m^-3^ d^-1^), is described as follows:

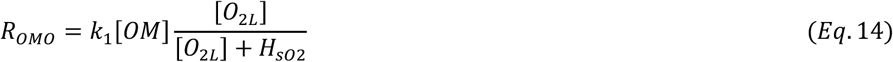

Total soil O_2_ includes O_2_ in dissolved form (*O_2L_*) (mol O_2_ per m^3^ soil water) and in gaseous form (*O_2G_*) (mol O_2_ per m^3^ soil void space). Soil O_2_ is influenced by O_2_ diffusion in gas form and in dissolved form, advection of dissolved O_2_, root respiration, oxidation of SOM and Fe(II). Dynamics of O_2_ are expressed as:

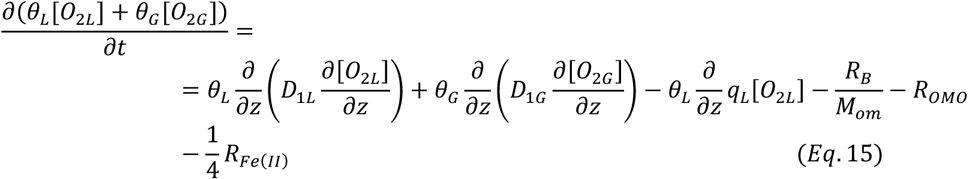

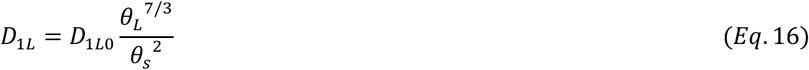

where *D*_1*L*_ and *D*_1*G*_ are diffusion coefficients (m^2^ s^-1^) of *O_2L_* and *O_2G_*. *D_1L_* varies with soil water content (*Eq*. 16). *θ_G_* represents soil gas content (m^3^ m^-3^). *θ_G_* changes in space and time, whose dynamics is modeled in the Section “*Soil Water Dynamics*” below. *D_1L0_* is the molecular diffusivity of O_2_ in the dissolved phase. At 25℃, [O_2L_] = 0.0318 [O_2G_], according to Henry’s law.

Clays, in addition to diffusion, are subject to disintegration and dispersion during Fe-C redox cycles, especially during Fe(III) reductive dissolution, when a large amount of colloids are markedly dispersed (37). We modeled clay changes with an empirical relationship determined at our study site, which shows that a decrease of 1 mol m^-3^ Fe(III) increases clay content by 0.0566% (Fig. S12). Similar negative associations between changes in clay content and in Fe(III) concentrations in regular redox patterns have been reported elsewhere (5).

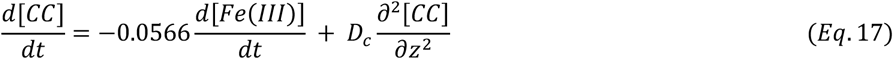

where *D_c_* is the clay diffusion coefficient (m^2^ s^-1^).

### Initial Conditions, Boundary Conditions, and Numerical Solutions

For model initial conditions, it is not feasible to obtain the state of system before the redox patterns emerged. We assume that soil conditions in our study area without regular redox patterns provide a good proxy for the initial condition. We used empirical measurements from nearby sites without redox patterns to initialize the model. Initial conditions for SOM are described by an exponential decay function of soil depth parameterized by field observations (Figs. S7 and S10):

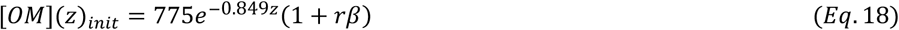

where *r* is a random number between −0.5 and 0.5 and *β* controls the magnitude of biomass fluctuations. We assume that OM at our study site consists of 5% living (root biomass, *B*) and 95% non-living parts (soil organic matter, *OM*)—that is, *B* is a linear function of *OM*. Initial Fe(II) was set to be zero everywhere in the model domain. Initial *O_2L_* is 0.273 mol m^-3^ everywhere in the soil profile, in equilibrium with the atmospheric O_2_. The initial condition of Fe(III) and clay concentrations follow an empirical hump shaped function with its peak at depth ∼ 1.2 m, informed by field measurements (Fig. S7). Fe(III) is at ∼ 45 mol m^-3^ in shallow soil layers (< 0.4 m), and between 438 and 500 mol m^-3^ in mid-layer soils (0.6 m ≤ depth < 1.5m). Between 0.4 and 0.6 m and below 1.5 m soil depth, Fe(III) is at the intermediate level (Fig. S7). In natural environments, the structure, solubility, and reactivity of Fe(III) minerals vary greatly (52, 86), and the same is true for OM (87). For the purpose of our study, our model describes general Fe chemical reactions in soils, without considering the heterogeneous reaction rates of diverse forms of Fe(III) and OM. As such, in the model, Fe(III) and OM can be depleted to zero via chemical reactions (*Eq*. 1).

For the upper boundary at the soil-air interface, we set a constant 0.273 mol m^-3^ for *O_2L_*, and a constant 0 for Fe(II), as we assume that Fe(II) is instantaneously oxidized by atmospheric O_2_. A constant of 1,162 g m^-3^ for root biomass density (*B*) was used for the upper boundary according to the empirical observation at the soil surface (4). The regional groundwater table is > 5 m and regular spatial patterns usually occur at soil depth < 2 m, below which are relatively homogenous Fe(III) distributions. As such, we set the lower boundary of the model at the soil depth of 4 m. We assume that root biomass is a constant zero at the lower boundary based on field observations (Fig. S10). For Fe(II) and *O_2L_*, we used a constant zero flux as the lower boundary. The model is solved by the implicit finite difference method, with the diffusion terms differentiated by the central difference scheme. An iterative method is used to address nonlinear model dynamics (79).

### Soil Water Dynamics

We modeled soil water flux (*q_L_*) and water content (*θ_L_*) dynamics by numerically solving the unsaturated flow equation—the mixed form Richards equation:

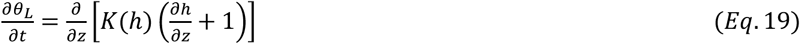

where *h* is pressure head (m), and *K* is hydraulic conductivity (m s^-1^). *K* is estimated following (88):

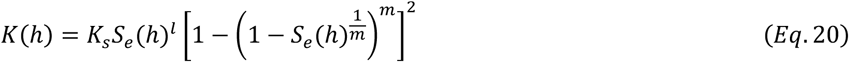

a

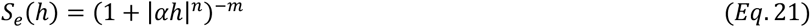

where *K_s_* is saturated hydraulic conductivity (m s^-1^), *l* is a pore connectivity parameter established to be 0.5 (89), *α*, *m* and *n* are semi-empirical fitting parameters of soil water retention curve (*m* = 1-1/*n*). *K_s_*, *α* and *n* are sensitive to soil texture. Redox oscillation modifies soil texture by its effect on clay disintegration, dispersion, and coagulation, thus soil texture, *K_s_*, *α* and *n* are dynamic in time and space. We allow *K_s_*, *α* and *n* to be a function of clay content (Supplementary Text S1), which evolves over time (*Eq*. 17); therefore, our model captures the evolution of soil texture over time, which directly affects soil water content and flux.

The Richards equation was solved using a mass-conservative finite difference method (90, 91). Due to the nonlinear nature of this equation, we employed Picard iteration and dynamic time step to ensure convergence of the solution at every time step. We applied daily rainfall and evapotranspiration observed between July 1, 2018 and July 1, 2019 at our study site as the climatic forcing (Fig. 7). While field measurements of O_2_, CO_2_, and soil volumetric water between 2016 and 2020 are available, the year 2018-2019 was chosen as the climatic forcing for the model because anoxic conditions were observed only during this year (Fig. S4). The same rainfall and evapotranspiration rates were repeated yearly in our model simulations, imposed as the dynamic upper boundary (soil-air interface). The upper boundary condition is either a prescribed flux or a prescribed head. By default, a prescribed flux equal to the observed evapotranspiration minus rainfall depth is used. However, when the calculated head (*h*) at the soil-air interface is > 0, the model assumes the existence of surface runoff during heavy rainfall periods, in which case, the upper boundary switches from the prescribed flux to a constant head of 0. We used free drainage (i.e., *∂h/∂z* = 0) for the lower boundary condition (at the soil depth of 4 m), and *h* = −1 m as the initial condition.

Solution of this model describes dynamic pressure head *h* of each computational grid. We used *Eqs.* 21 and 22 to calculate dynamic *S_e_* and *θ_L_* respectively. Darcy’s law was used to estimate spatially and temporally varying *q_L_*:

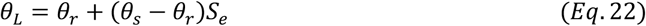

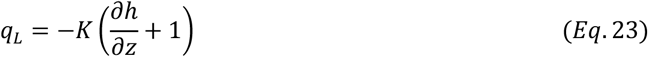

Where *θ_r_* is residual water content of soils. *θ_r_* and *θ_s_* vary with soil texture—in our study, the temporal evolution of clay content (Supplementary Text S1).

### Numerical Experiments and Model Analysis

To distinguish among alternative hypotheses of pattern formation, we tested the following model variants (Table S3). (1) To test the mechanism of coupled amplifying SDF with threshold dependent SDF based on root responses to Fe(III), we ran the full baseline model described above. (2) To test whether the amplifying SDF realized by clay dynamics *alone* can generate regular patterns, we switched off processes associated with SDF based on root responses to Fe(III), by setting *f_Fe(III)_* in *Eq*. 10 to be 1 (Table S3); that is, plant growth is no longer affected by Fe(III). (3) To test whether the threshold dependent SDF based on root responses to Fe(III) *alone* can produce regular patterns, we switched off the amplifying SDF by removing the term *Se^φ^* in *Eq.* 4 (Table S3); (4) To investigate the mechanism of monotonic SDF based on root responses to water limitation, we turned off the amplifying SDF and threshold dependent SDF. It is realized by setting *f_Fe(III)_* in *Eq*. 10 to be 1 and removing the term *Se^φ^* in *Eq.* 4. The effect of soil water content on root growth is incorporated in the root growth equation by adding *Se^2^* to *Eq*. 10 (Table S3). Lastly, (5) to test the hypothesis that the observed redox pattern is caused by a preexisting regularly distributed root structure, we prescribed regularly distributed root biomass and SOM as model initial conditions (Fig. S2). Meanwhile, the carrying capacity *K_B_* is set to equal the initial-condition distribution of SOM (Table S3). This allows the root biomass to maintain the same regular patterning over time. All SDFs were switched off in the model and the root diffusion term was removed from the biomass change function (*Eq*. 8; Table S3). Patterns—including vertical concentration distributions of Fe(III) and OM (biomass *B* is a linear function of OM) —generated by different model variants are compared with the patterns observed in the field to determine the most plausible mechanism.

We investigated the effect of climatic, soil, and biological variables on emergence of regular patterns, characteristics of patterning (e.g., patterning range, banding width), and the time it required for patterns to form. Pattern formation time is defined as the time it requires for the patterns to statistically stabilize *and* for Fe(III) to reach 138 mol m^-3^, the observed Fe(III) concentration in gray layers (Table S2) at our study site (4). To test the effect of climatic condition and soil texture, we manipulated annual precipitation regime and soil clay content, respectively (Supplementary Text S2). The effect of diffusion rates on the banding width is tested by varying *D_b_* (biomass diffusion; *Eq*. 8) and *D_1_* (Fe(II) diffusion; *Eq*. 7).

### Estimating the Effect of Pattern Formation on Soil Carbon Storage

We assessed consequences of redox pattern formation on soil carbon storage. We first digitized photos of empirical patterns in the pattern formation zone of our study site (Fig. 1D) to quantify the width of gray and of orange layers. We then calculated the total carbon storage in the pattern formation zone (between 1.0 m and 1.8 m) by multiplying the width by measured layer-specific SOC content. This calculated carbon storage value was compared with the carbon storage in the same zone from a nearby site without regular patterning. This comparison allows us to infer the percent reduction in carbon storage capacity attributable to pattern formation. Furthermore, with the model, we compared the difference of total carbon in the initial condition and in the steady state condition after the pattern is formed. This allows us to calculate the percent change in soil carbon storage over the period of pattern formation. Since we used the SOC profile measured at the site without patterning as the model initial condition, we expect that the percent change of carbon storage from the initial condition to the steady state to be the statistically similar to the percent difference between the sites with and without patterning.

### Evaluating Model Performance

The baseline model captured the overall vertical profiles of Fe(III) and OM, as well as O_2_ profiles during relatively dry periods of soil oxic conditions and during wet, anoxic conditions (Fig. 4). In addition to the overall profiles, the model reproduced the distinctive concentration contrast of Fe(III), OM, root biomass, clay, and soil water content between gray and orange layers (Fig. 4). The model slightly overestimated Fe(III) in orange layers and underestimated OM there (Fig. 4; Table S2). This is likely because with the increase of Fe(III) in orange layers, the diffusion coefficient of Fe(II) can become much smaller, but such an effect was not fully incorporated in our model, due to lack of experimental data. Furthermore, the model faithfully reproduced time-series of O_2_ and soil volumetric water content (Fig. S5). Finally, the model reproduced spatial patterns that are statistically the same as observed patterns, including both the spatial extent (depth of upper and lower boundaries of patterned section), width of orange and gray layers, and their spatial regularity (Fig. 4).

## Classification

Physical Science + Earth, Atmospheric, and Planetary Science Biological Science + Ecology

## Acknowledgement

We thank Stuart G. Fisher for discussions that significantly improved this paper.

---

**Table S1.**
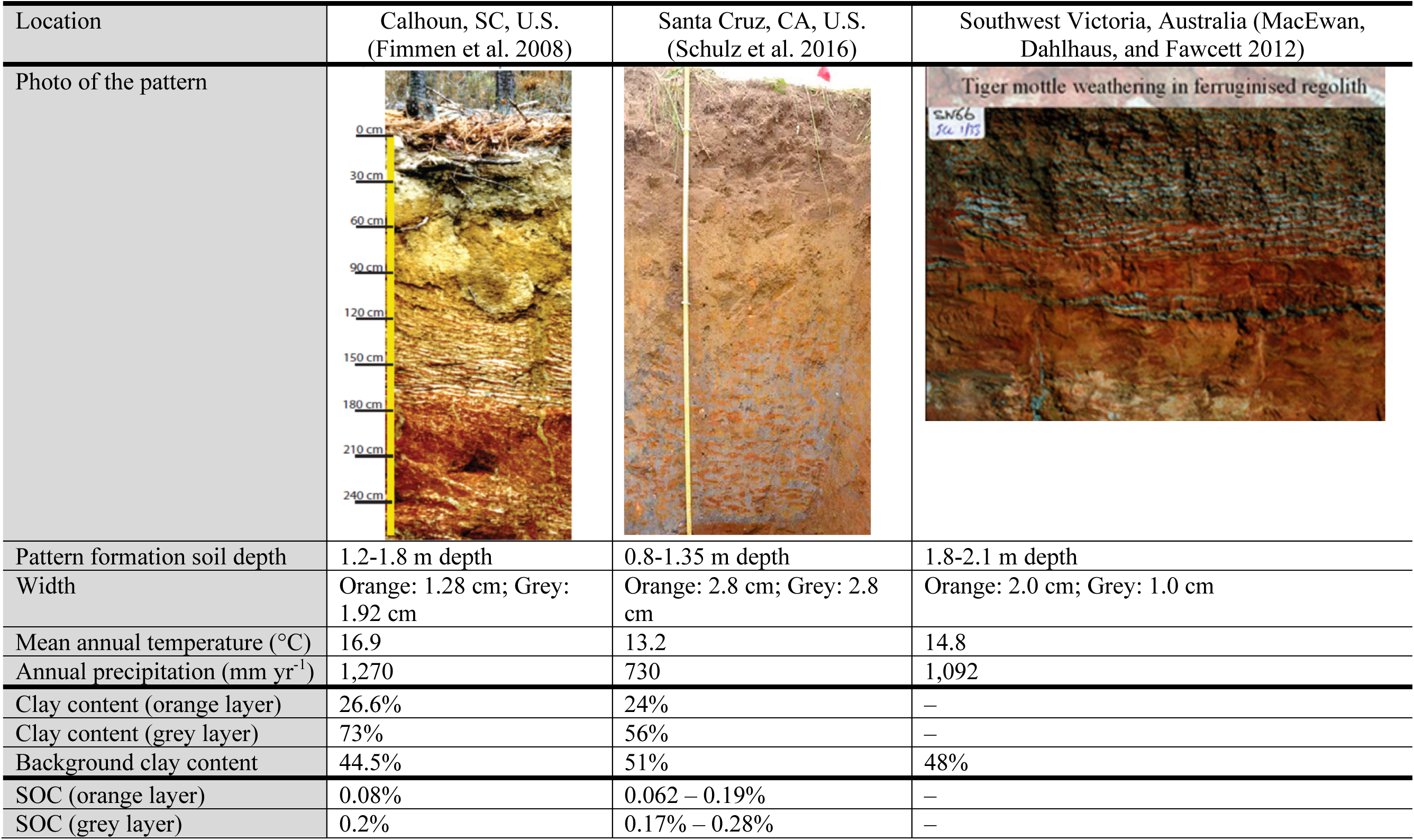
Climatic and soil conditions of sites where regular iron redox patterns in soils have been reported. Note that we only collected papers that explicitly describe that the redox pattern is regular or show images of regular patterns.

**Table S2.**
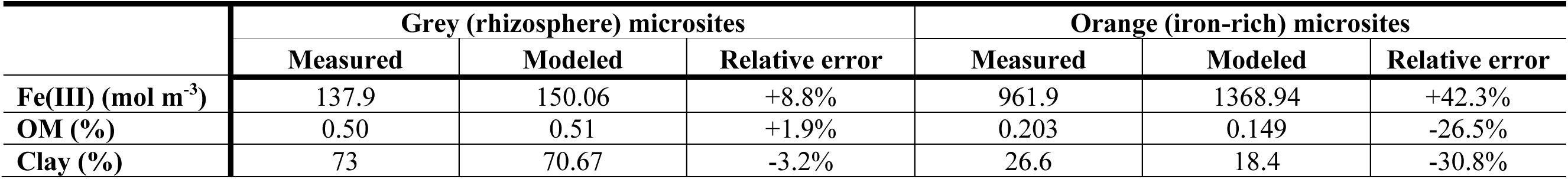
Comparison between the measured and modeled Fe(III) concentration, OM content, and clay content in the gray layers and in the orange layers of the formed pattern at our study site.

**Table S3.**
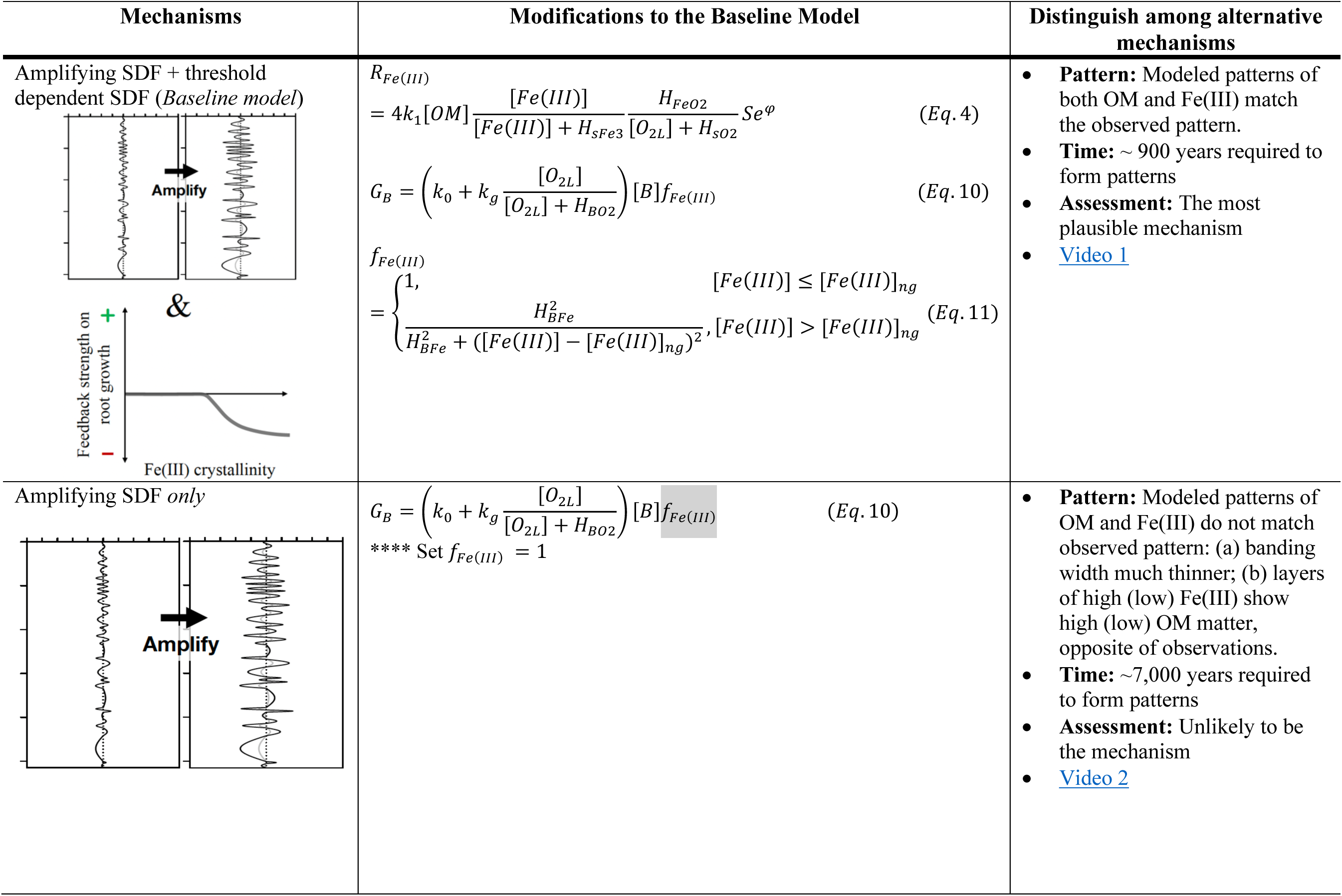
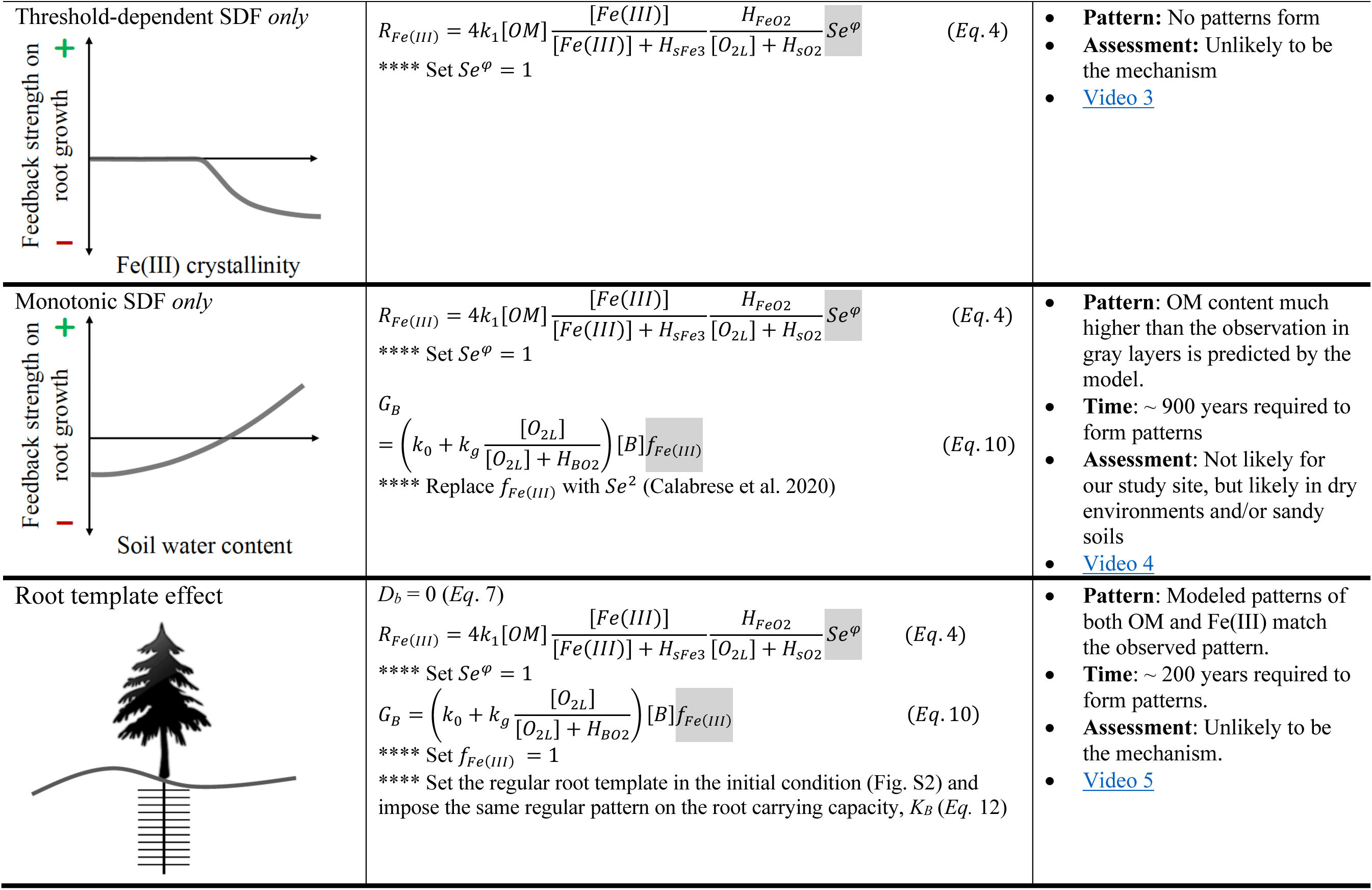
The major hypothesized mechanisms, their corresponding model representations, and differences in their model predictions of redox patterns. The terms that are modified for each hypothesis are highlighted in gray.

**Table S4.**
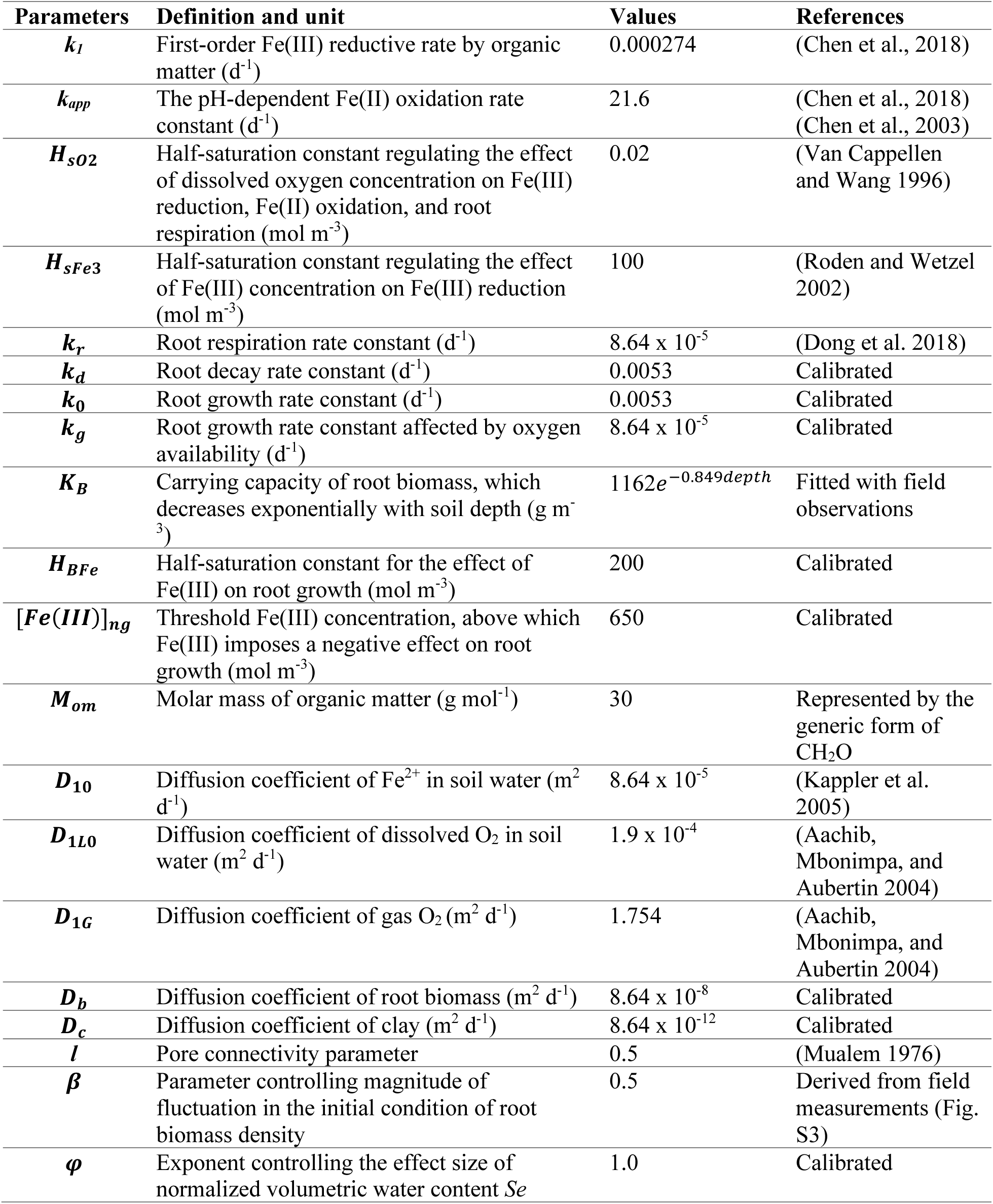
Definition, notations, and values of model parameters.

Video 1: Formation of redox patterns in upland soils by the mechanism coupling threshold-dependent SDF and amplifying SDF. The first row of plots shows profiles of the various chemical concentrations in the entire model domain (0-to-4-meter soil depth), including—from left to right—dissolved O2 concentration (%), dissolved Fe(II) (mol m^-3^), Fe(III) deposit (mol m^-3^), and organic matter (mol m^-3^). The second row of plots shows the pattern in the zone between 1.2 and 1.5 m soil depth. From left to right: soil volumetric water content, concentration of Fe(III) deposit (mol m^-3^), clay content (%), and organic matter content (mol m^-3^). The simulation shows a time interval of 1 year over ∼ 800 years.

Video 2: Formation of redox patterns in the upland soils by the mechanism of amplifying SDF alone. The first row of plots shows profiles of the various chemical concentrations in the entire model domain (0-to-4-meter soil depth), including—from left to right—dissolved O_2_ concentration (%), dissolved Fe(II) (mol m^-3^), Fe(III) deposit (mol m^-3^), and organic matter (mol m^-3^). The second row of plots shows the pattern in the zone between 1.2 and 1.5 m soil depth. From left to right: soil volumetric water content, concentration of Fe(III) deposit (mol m^-3^), clay content (%), and organic matter content (mol m^-3^). The simulation shows a time interval of 10 year over ∼ 4,000 years.

Video 3: Formation of redox patterns in the upland soils by the mechanism of threshold-dependent SDF alone. The first row of plots shows profiles of the various chemical concentrations in the entire model domain (0-to-4-meter soil depth), including—from left to right—dissolved O_2_ concentration (%), dissolved Fe(II) (mol m^-3^), Fe(III) deposit (mol m^-3^), and organic matter (mol m^-3^). The second row of plots shows the pattern in the zone between 1.2 and 1.5 m soil depth. From left to right: soil volumetric water content, concentration of Fe(III) deposit (mol m^-3^), clay content (%), and organic matter content (mol m^-3^). The simulation shows a time interval of 1 year over ∼ 1,000 years.

Video 4: Formation of redox patterns in the upland soils by the mechanism of monotonic SDF. The first row of plots shows profiles of the various chemical concentrations in the entire model domain (0- to-4-meter soil depth), including—from left to right—dissolved O_2_ concentration (%), dissolved Fe(II) (mol m^-3^), Fe(III) deposit (mol m^-3^), and organic matter (mol m^-3^). The second row of plots shows the pattern in the zone between 1.2 and 1.5 m soil depth. From left to right: soil volumetric water content, concentration of Fe(III) deposit (mol m^-3^), clay content (%), and organic matter content (mol m^-3^). The simulation shows a time interval of 1 year over ∼ 1,000 years.

Video 5: Formation of redox patterns in the upland soils by the mechanism of pre-existing root template. The first row of plots shows profiles of the various chemical concentrations in the entire model domain (0-to-4-meter soil depth), including—from left to right—dissolved O_2_ concentration (%), dissolved Fe(II) (mol m^-3^), Fe(III) deposit (mol m^-3^), and organic matter (mol m^-3^). The second row of plots shows the pattern in the zone between 1.2 and 1.5 m soil depth. From left to right: soil volumetric water content, concentration of Fe(III) deposit (mol m^-3^), clay content (%), and organic matter content (mol m^-3^). The simulation shows a time interval of 1 year over 180 years.

**Figure S1.**
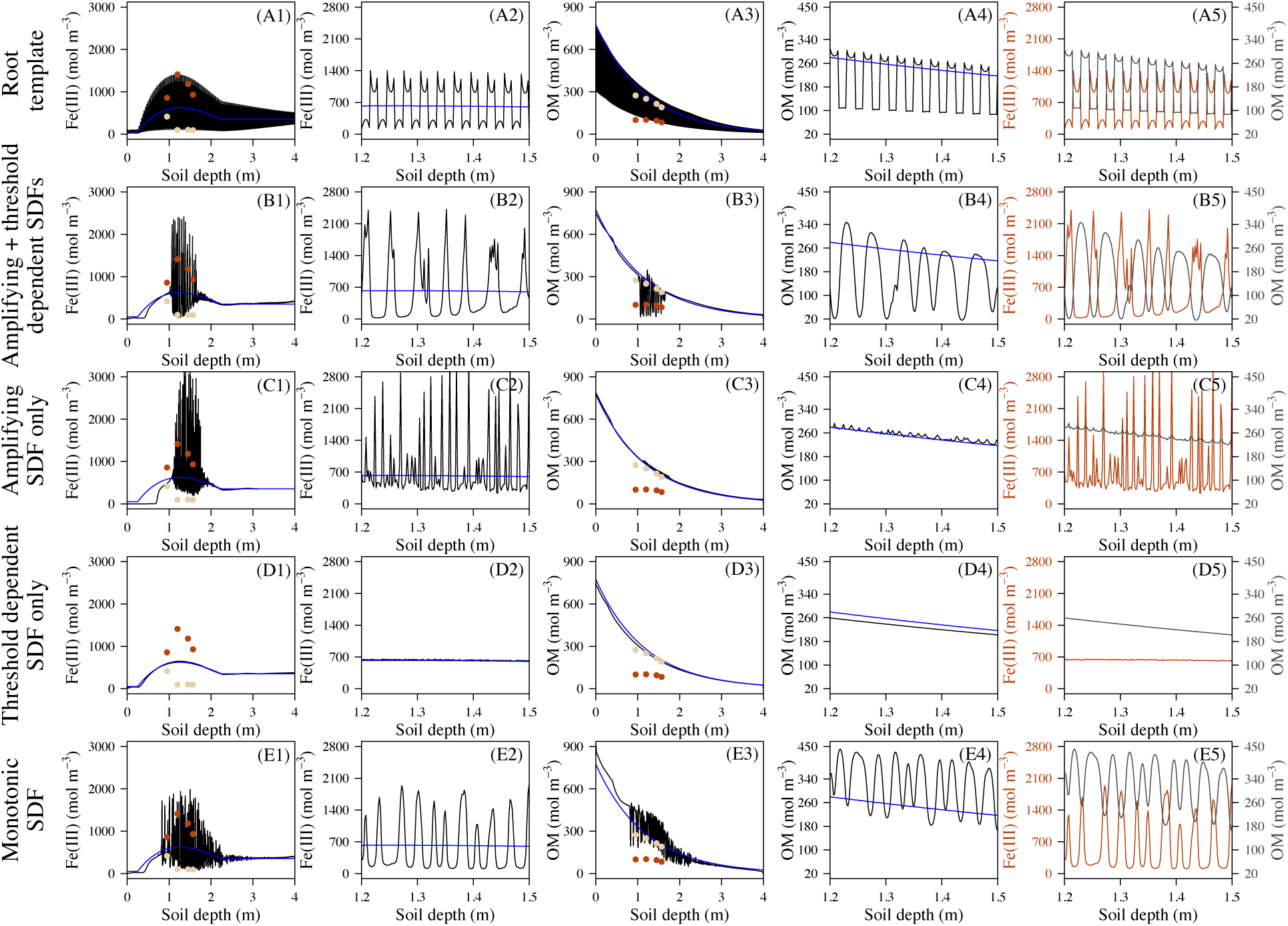
Model simulated patterns with five hypothesized mechanisms: Patterns formed by root template effect (A1-5); by coupling the amplifying SDF with threshold dependent SDF based on negative root responses to Fe(III) (B1-5); by the amplifying SDF alone (C1-5); by the threshold dependent SDF based on negative root responses to Fe(III) alone (D1-5); and by the monotonic SDF based on root responses to soil water content (E1-5). Blue lines in each plot show the observed background concentrations, represented by the data from nearby sites without patterning. Red points are empirical measurements from the Fe(III) concentrated orange layers at our study site, and the light-yellow points are empirical measurements from the Fe(III) depleted gray layers. The first column describes the modeled Fe(III); the second column is a close-up view of the Fe(III) in the pattern formation zone between 1.2 and 1.5 m soil depth; the third column describes the modeled organic matter (OM); the fourth column is a close-up view of OM in the pattern formation zone; and the last one is overlapping Fe(III) and OM profiles to show their spatial relationships of high and low.

**Figure S2.**
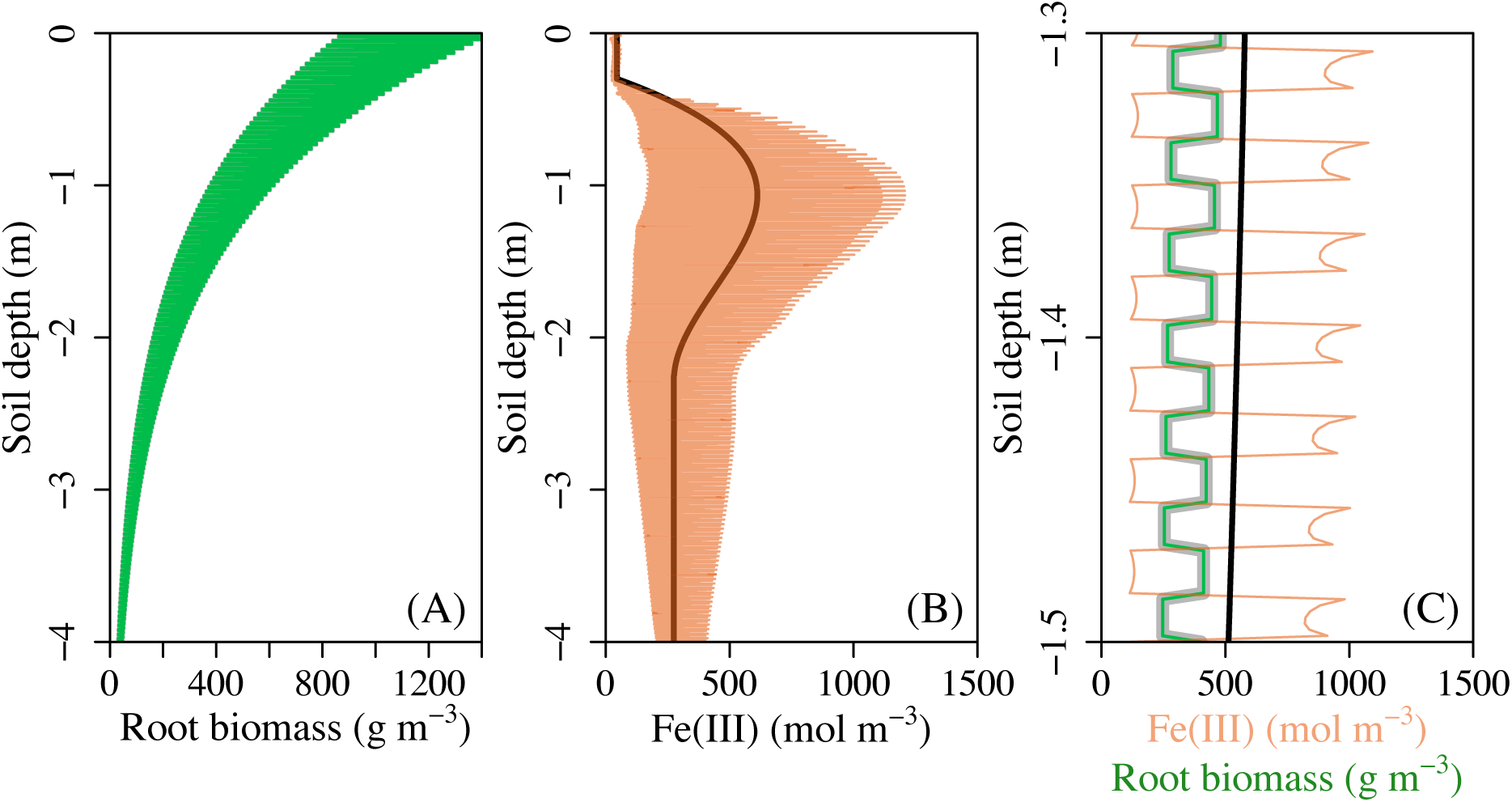
Redox patterns follow the *exact* pattern set by the root biomass in the initial condition. (A) shows the root biomass in the steady state (green) and in the initial condition (gray; as it overlaps exactly with steady-state distribution, gray line is not visible); (B) shows the Fe(III) profile in the steady state (orange) and in the initial condition (black); and (C) zooms in to the soil depth between 1.3 m and 1.5 m showing that root biomass in the initial condition (thick gray line) and in the steady state (green) and the initial-state (thick black line) and steady-state (orange) Fe(III) concentration distribution.

**Figure S3.**
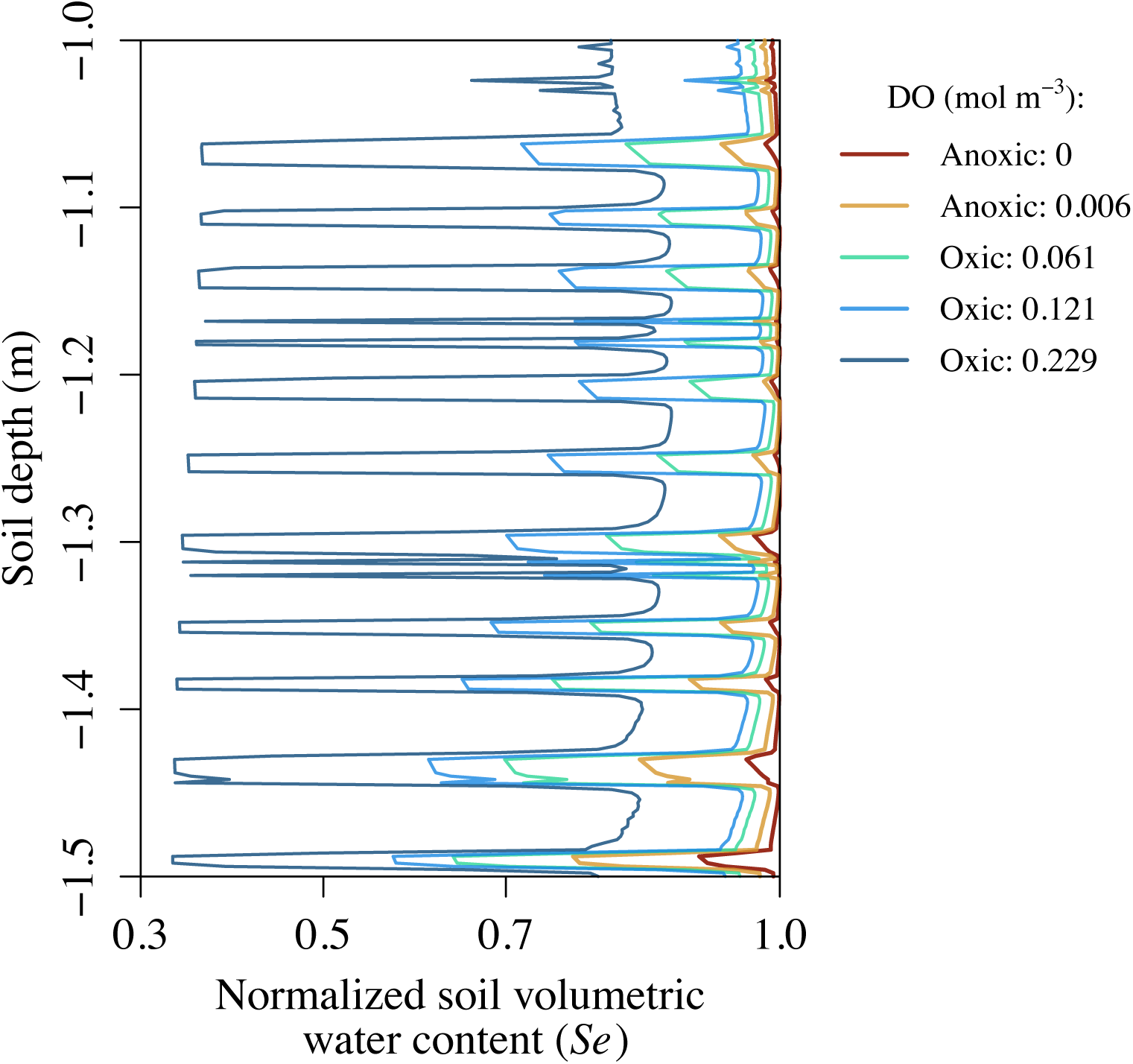
Difference of soil water content between orange and grey layers along the anoxic-oxic gradient, represented by the modeled soil water content in the pattern formation zone between 1.0 and 1.5 m soil depth. When soil is under anoxic condition, the difference of water soil content between orange and gray layers is small, resulting in weak amplifying effect. As soil gets drier and becomes more oxic, the soil water content between orange and gray layers becomes large, allowing a relatively strong amplifying effect occur. However, under oxic conditions, reductive dissolution of Fe(III) is suppressed. As a result, the amplifying effect is again limited. DO concentrations represent the average concentration between the soil depths of 1.0 and 1.5 m.

**Figure S4.**
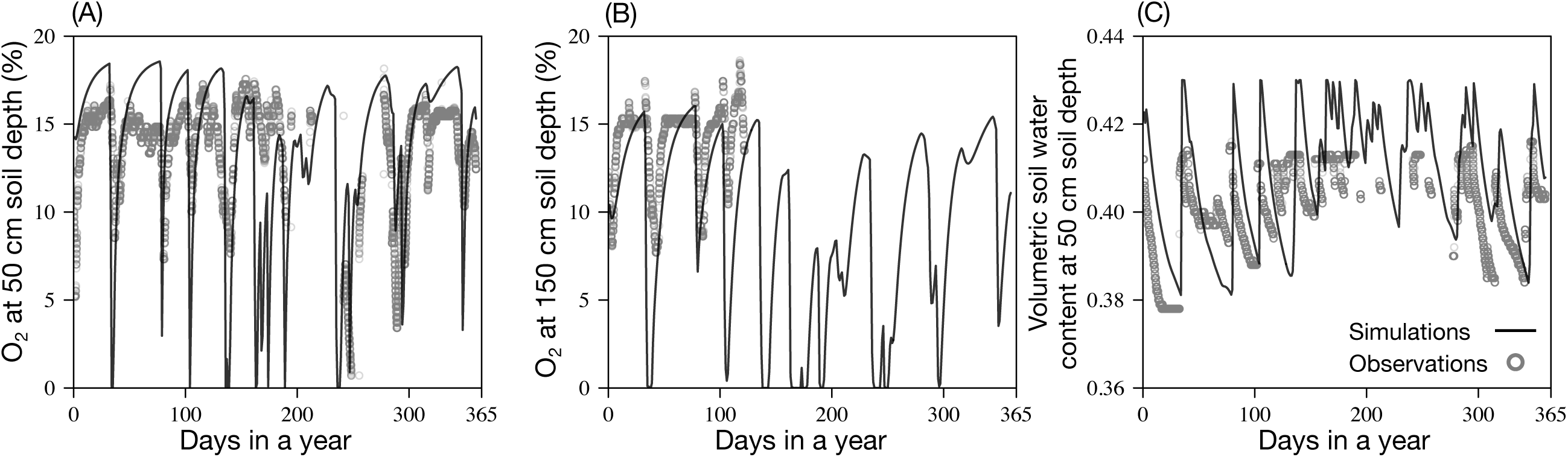
Comparison of the observed (between July 1, 2018 and July 1, 2019) and modeled O_2_ concentrations at 50 cm (A) and 150 cm (B) soil depth and soil volumetric water content (C). Our model provides a good fit to the empirical data.

**Figure S5.**
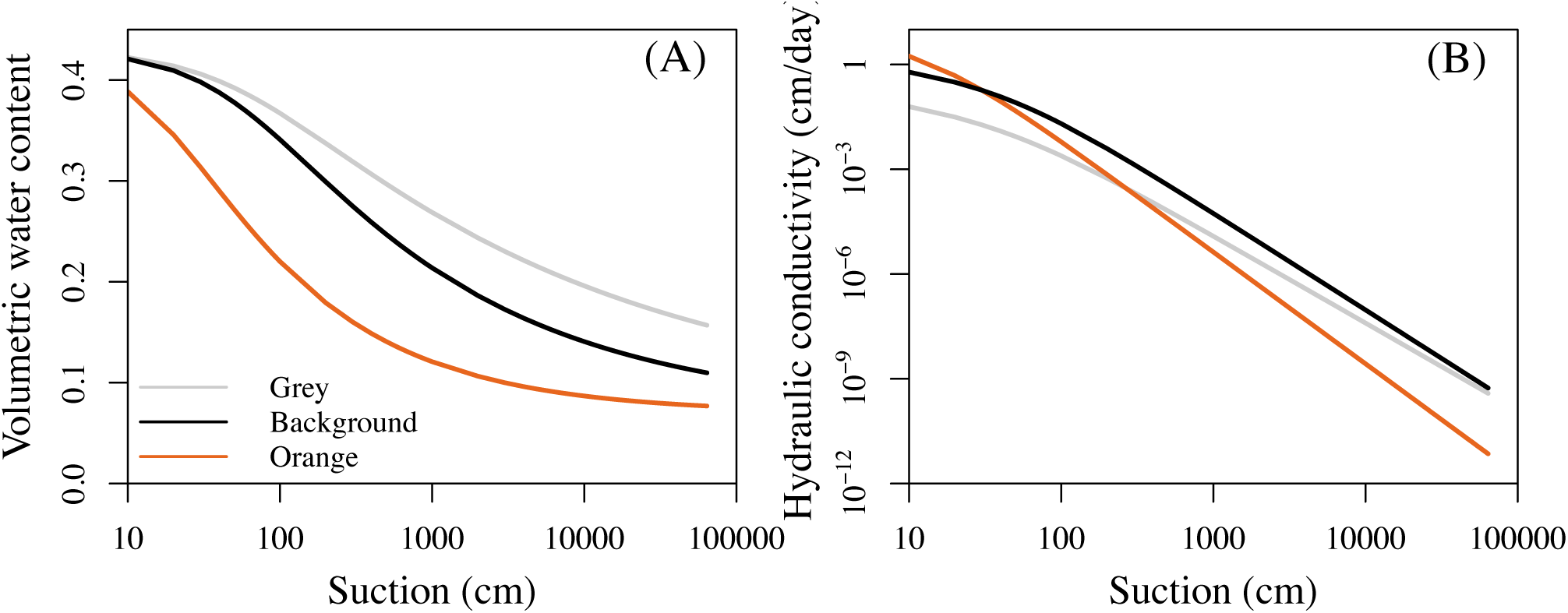
Modeled water retention curve and hydraulic conductivity curve for the soil texture in the background condition and for the contrasting soil textures in gray layers and orange microsites in regular redox patterns (Table S5 in Supplementary Text provides descriptions of characteristics of these three types of soil texture). Suction = 0 – pressure head, *h*.

**Figure S6.**
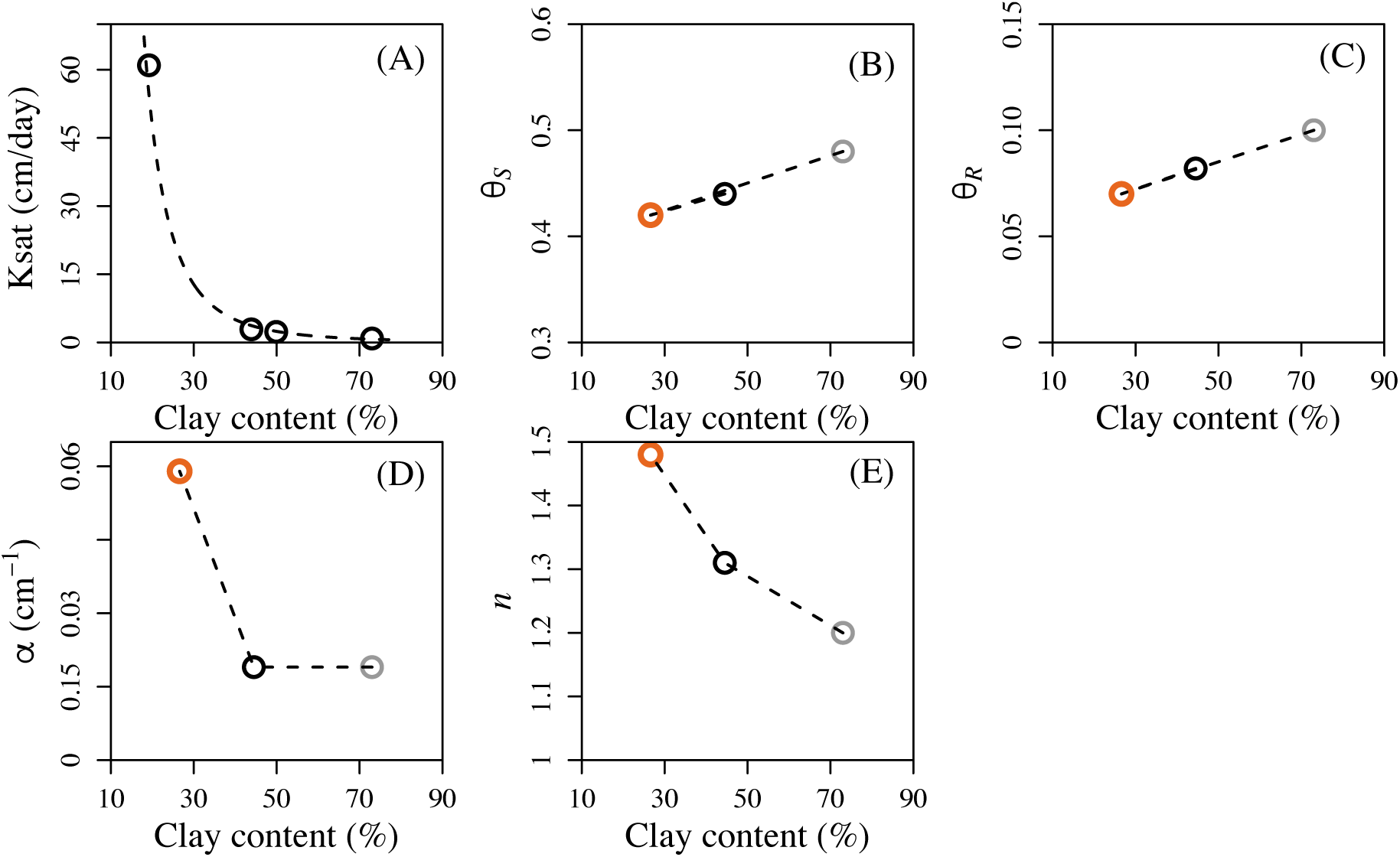
Empirical relationships between five hydraulic parameters (Table S5; Text S1) and clay content used in the model: (A) saturated hydraulic conductivity, *K_s_*, (B) saturated soil water content, *θ_s_*, (C) residual soil water content, *θ_r_*, (D) fitting parameter, *α* (*Eq*. 19) (E) fitting parameter, *n* (*Eq*. 19). In (B)-(E): orange circles represent samples from orange layers of redox patterns at our study site, gray circles represent samples from gray layers, and black circles represent samples from bulk samples, as background level. Piecewise linear regressions were used to represent the relationship between the four hydraulic parameters and clay content (B-E), and a fitted exponential relationship, *K_s_* = 807241x (clay content)^-3.24991^, was used for the relationship between *K_sat_* and clay content (A). These empirical relationships were used to model soil water dynamics.

**Figure S7.**
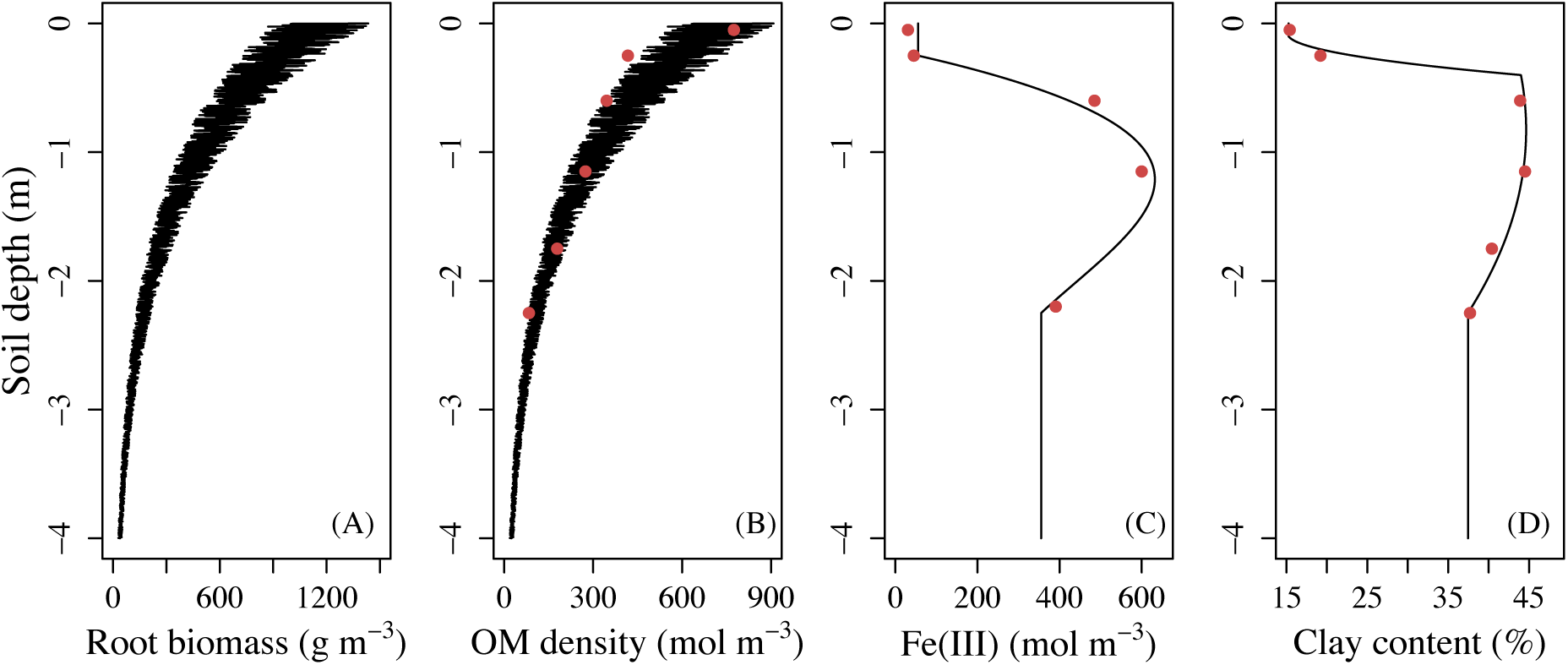
Initial conditions used in the model for (A) soil vertical profiles of root biomass, (B) organic matter concentration (OM), (C) Fe(III) concentration, and (D) clay content. The initial conditions are informed by observed patterns (red points) at nearby sites without patterns.

**Figure S8.**
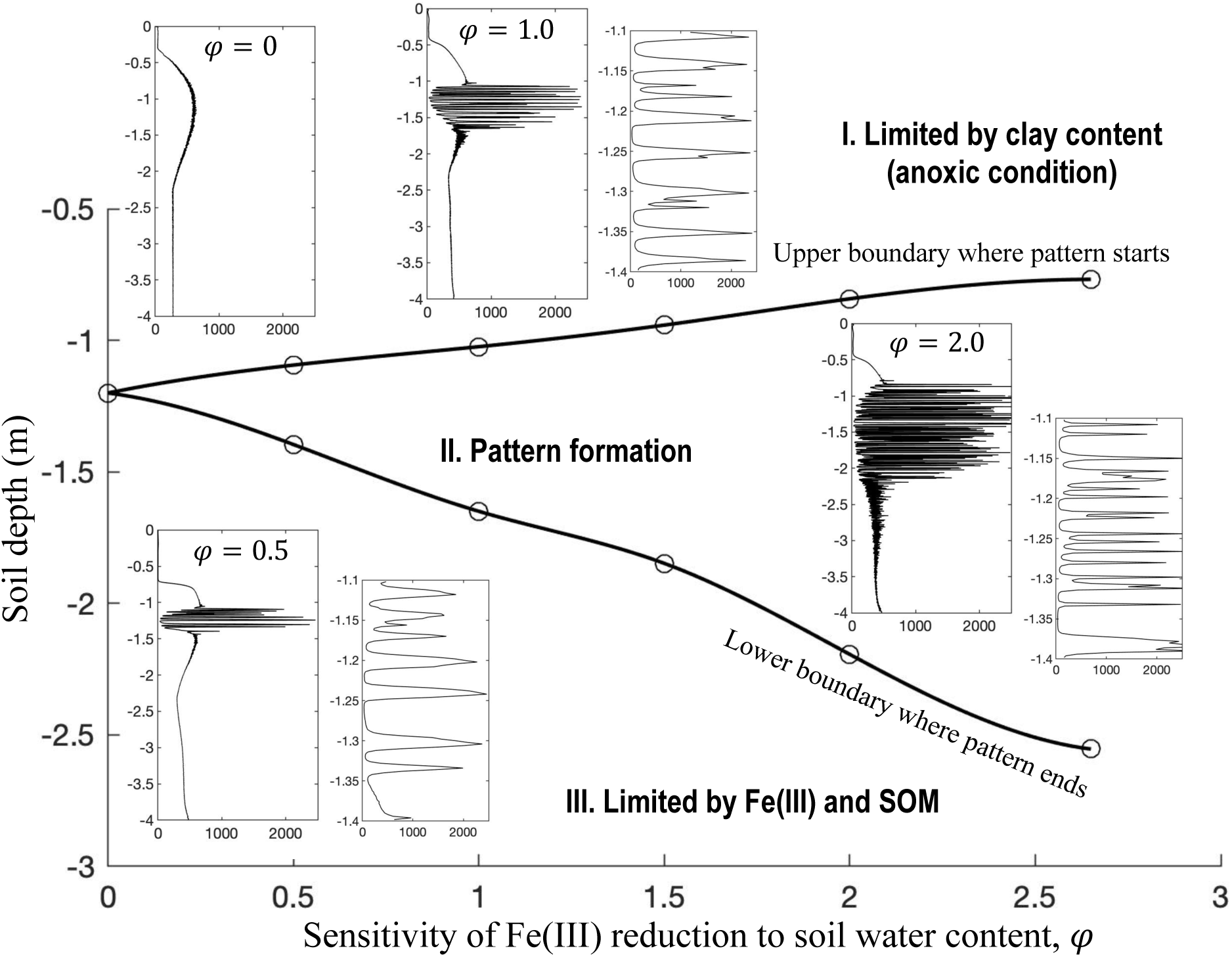
Sensitivity of Fe(III) reduction rate to soil water content (*φ* in *Eq*. 4) affecting the spatial extent of pattern formation zone in the soil profile.

**Figure S9.**
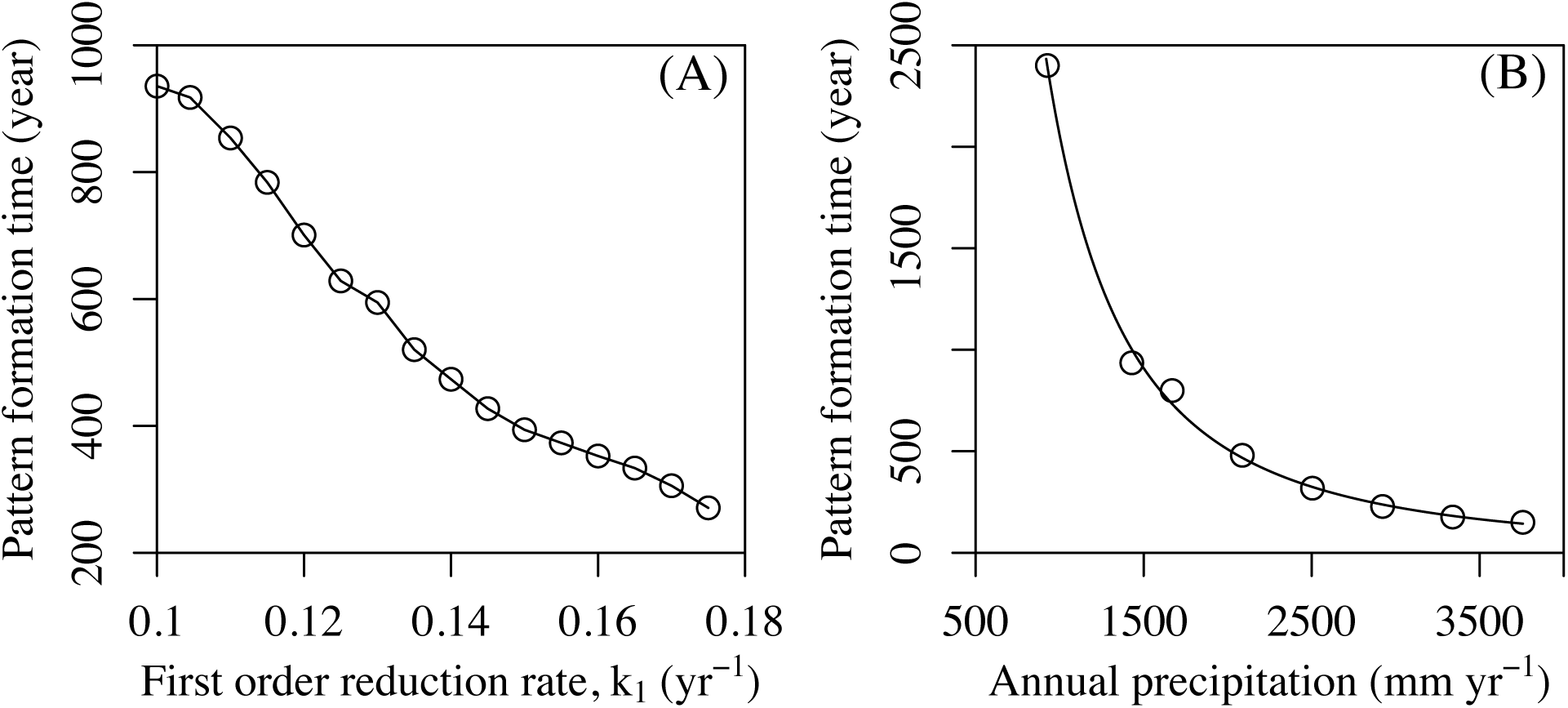
Time required for redox pattern formation in upland soils affected by first order Fe(III) reduction coefficient by soil organic matter, *k_1_* (A) and by annual precipitation (B). Pattern formation time changes linearly with *k_1_*, but exponentially with precipitation (*R^2^*> 0.99).

**Figure S10.**
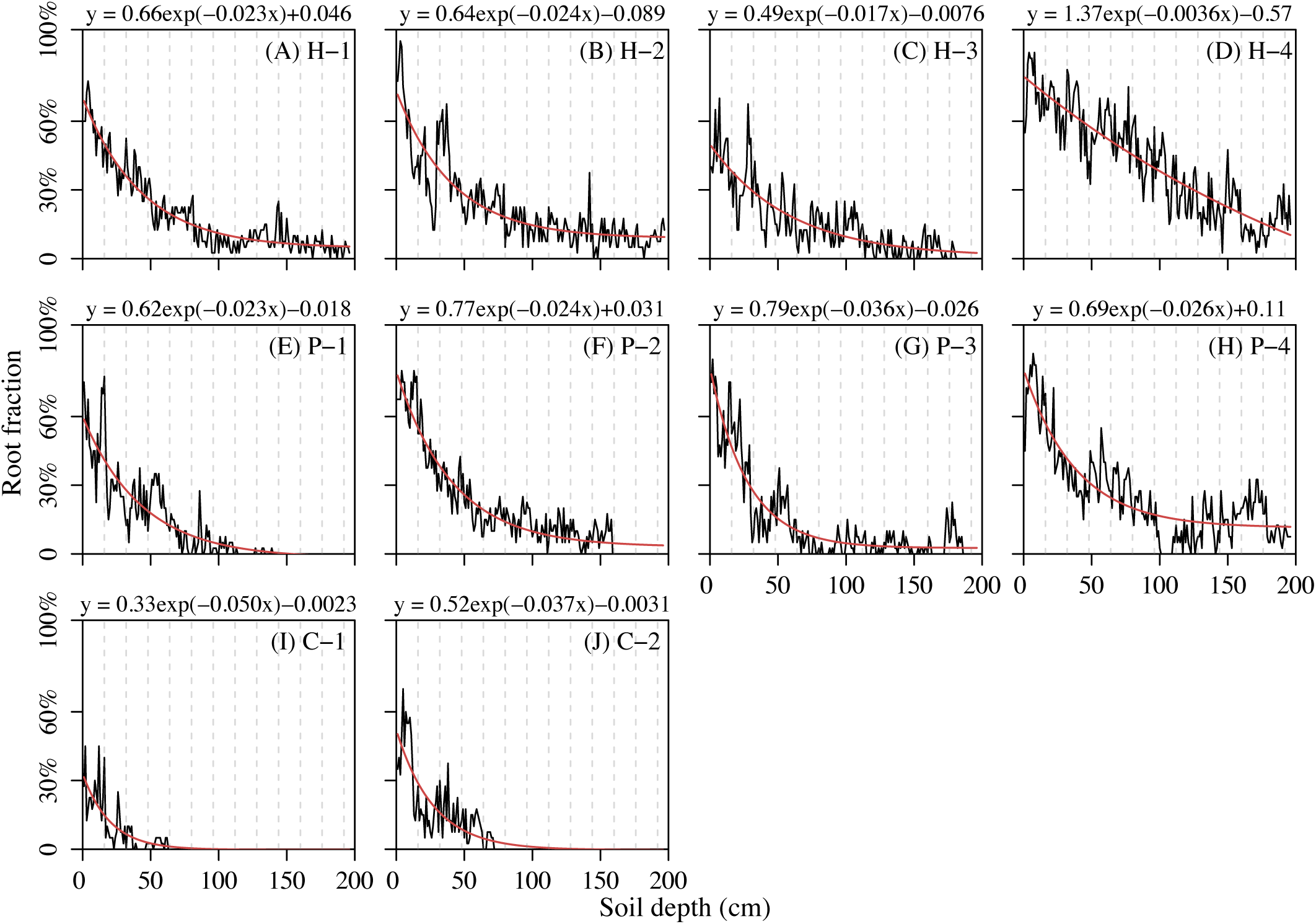
Measured (at every 1 cm) fine root distribution at ten different sites of the Calhoun Experimental Forest (South Carolina, U.S.) with best fit exponential decay functions. Dominant plant species varies among sites: hardwood trees (“*H*”), pine trees (“*P*”), and cottons (“*C*”). Biomass of cotton roots declines more rapidly than that of hardwood trees or pine trees. The dashed lines in each plot are 1.6 cm apart, the average width of an orange or gray layer in the regular redox patterns in our study site. Roots do not show regularly spaced (at 1.6 cm interval) biomass distribution (as in Figure S2).

**Figure S11.**
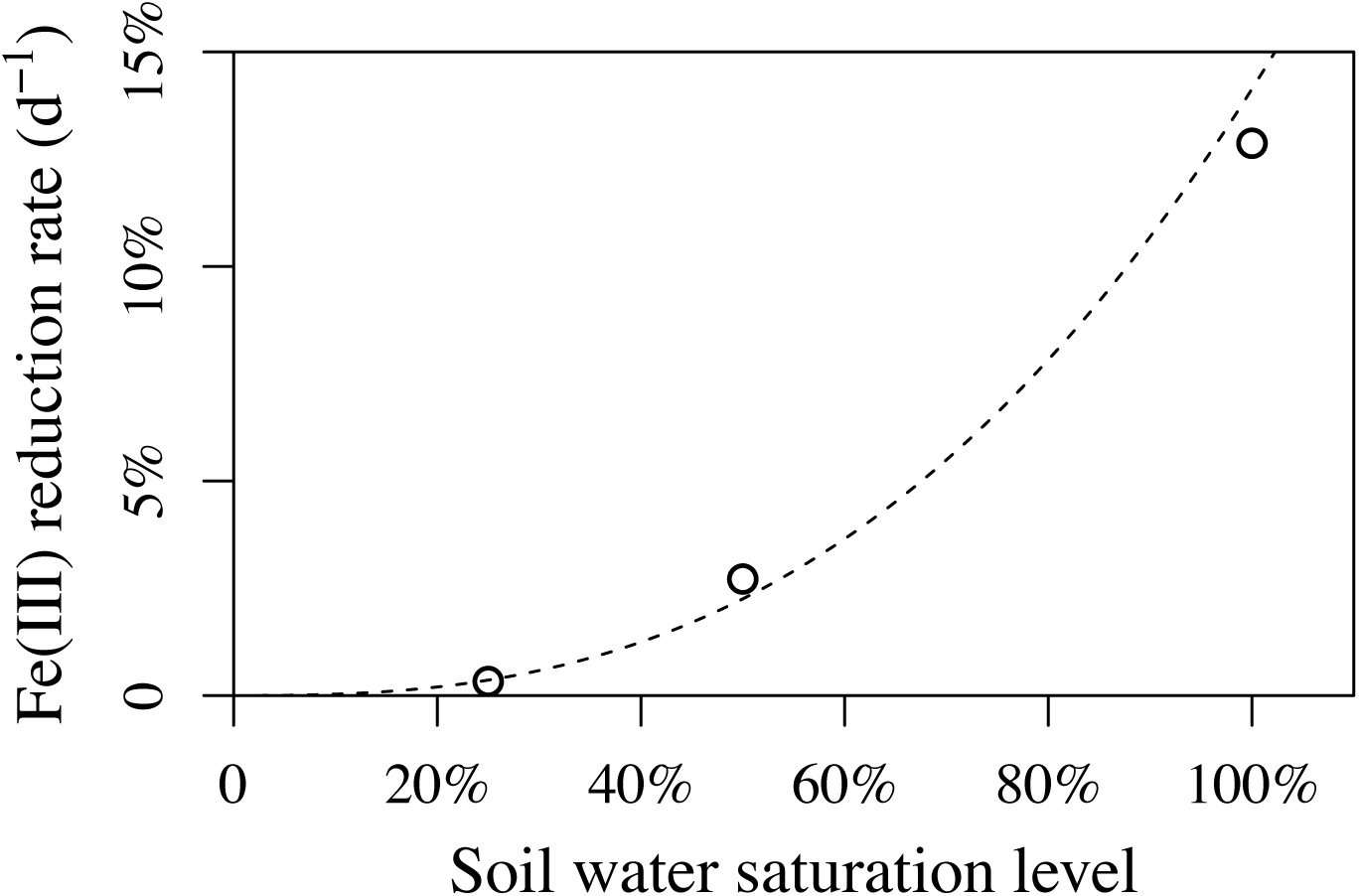
The effect of soil water content on the rate of Fe(III) reduction. The data are extracted from Figure 2 of the paper by Hodges et al. (2018). Fe(III) reduction rate represents the fraction of Fe(III) removed by reductive dissolution after two weeks of lab experiments, controlled at different levels of soil water saturation.

**Figure S12.**
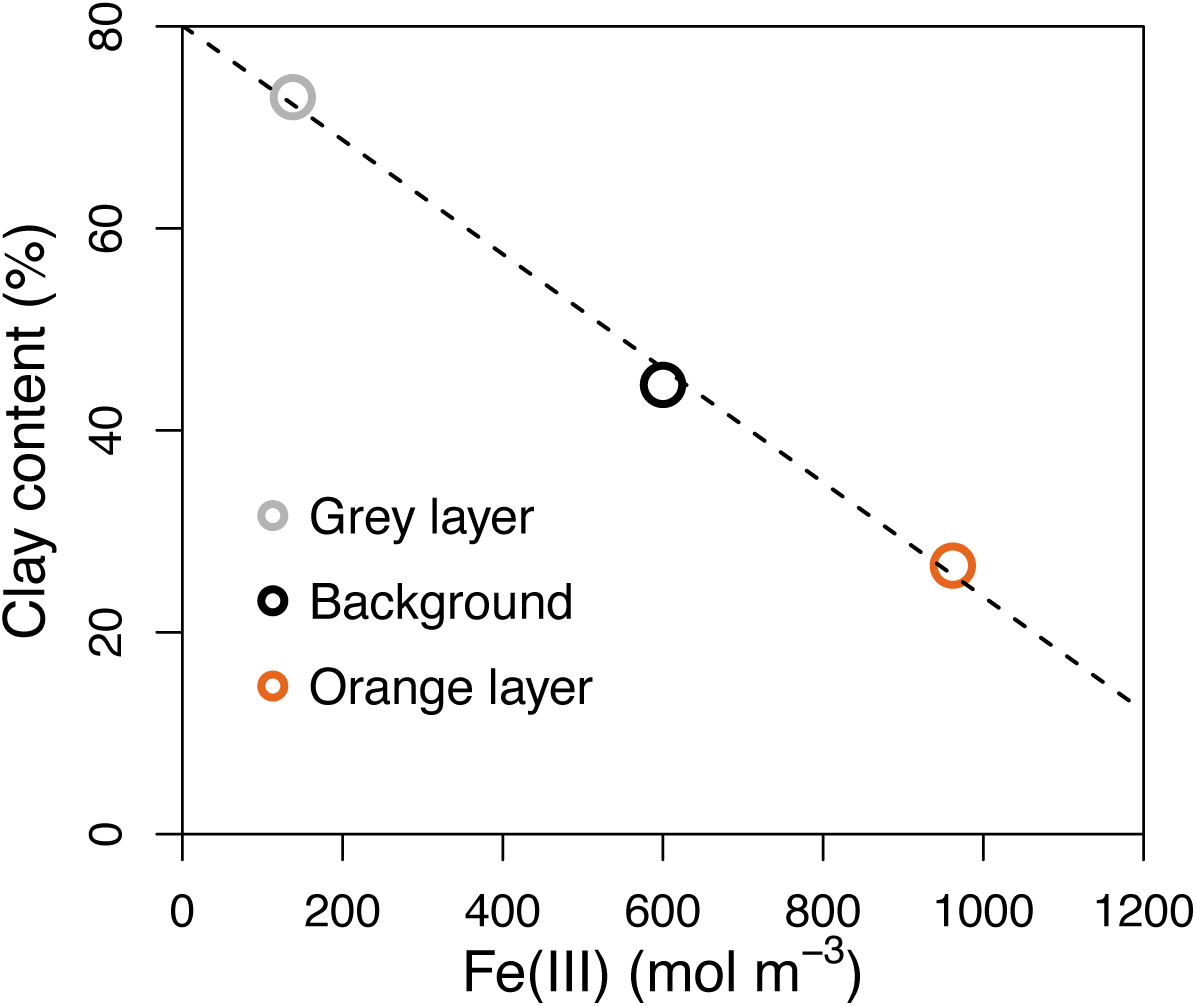
Empirical relationship between Fe(III) concentration and clay content with data from the Calhoun Experimental Forest (South Carolina, U.S.). This suggests the effects of iron redox reactions on clay disintegration and coagulation. Different colors represent the source of the data: Data from orange and gray layers in the regular redox patterns are in orange and gray circles, respectively; and data from sites without redox patterns, representing the background condition, are in black.

## Supplementary Text S1

### Relationships between soil hydraulic parameters and clay content in soils

The progression of iron redox reactions continuously modifies clay content in soils, hence changing soil texture. Soil texture has a significant effect on water content and water flux, which in turn feeds back to affect soil redox dynamics and iron chemical reactions. To capture such feedbacks, we modeled the dynamics of soil hydraulic parameters based on empirical measurements in our study site and model calibration. We considered the impact of clay concentration on the five hydraulic parameters in *Eqs.* 19-21: saturated hydraulic conductivity (*K_s_*), fitting parameters *α* and *n*, saturated water content (*θ_s_*), and residual water content (*θ_r_*). As clay concentration increases, soil texture becomes finer, water permeability decreases, i.e., lower *K_s_*, *α* and *n*, and water retention capacity increases, i.e., higher *θ_s_* and *θ_r_* (Simunek, Van Genuchten, and Sejna 2005; Brogowski, Kwasowski, and Madyniak 2014). To represent these dynamics in the model, we compiled the soil texture data measured at gray layers and orange layers in the pattern formation zone in our study site. Since it is not feasible to have reliable information on soil texture in the initial condition before the regular redox patterns were formed, we used soil texture data collected at a nearby site without regular patterns (defined as “background” level in this study) to represent the soil texture in the initial condition. The soil texture data and hydraulic parameters were extracted from (Fimmen et al. 2008) and shown in Table S5.

**Table S5.**
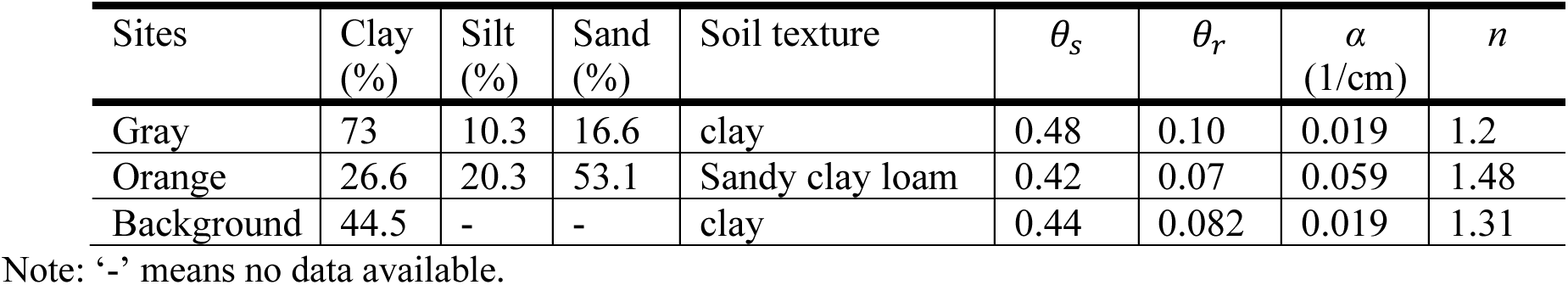
Soil texture and hydraulic parameters for the grey and the orange layers and proxy of the initial condition with data collected from a nearby site without redox patterning (“*background*”).

When percent of clay is known, *θ_s_* is estimated following the method by (Brogowski, Kwasowski, and Madyniak 2014). We inferred values of *θ_r_*, *α*, and *n* using Rosetta, a computer program to estimate soil hydraulic parameters for a given soil texture (Schaap, Leij, and Van Genuchten 2001). The four hydraulic parameters at a given clay concentration were estimated using a piece-wise linear interpolation (Vogel, Cislerova, and Hopmans 1991) (Fig. S11B-E). *K_s_* values for soils of different clay contents were directly measured in the field. Based on the observed *K_s_* and clay content relationship, we fitted an exponential relationship (Fig. S11A). This fitted function is used in the model to inform *K_s_* under any given clay content. Our model simulated spatially and temporally varying clay content (*Eq.* 17). At each step, with the evolution of clay content, the values of these five parameters also change as a function of clay content according to the relationships in Fig. S11. Taking these steps, the model reproduced soil water content and oxygen dynamics reasonably well with field observations (Fig. S5).

## Supplementary Text S2

### Manipulate precipitation regime and soil clay content in numerical experiments

To test the effect of precipitation regime on pattern formation, we first downloaded daily precipitation and evaporation rate between 1950 and 2021 at the weather station in the Union County, S.C. (U.S.A.) from National Centers for Environmental Information at NOAA (https://www.ncei.noaa.gov/cdo-web/datasets). We picked four typical years representing wet (1971), normal (1990), dry (1952), and very dry (2007) conditions with an annual precipitation of 1,669.8 (100% quantile), 1,313.2 (74% quantile), 927.1 (20% quantile), and 528.6 mm yr^-1^ (20% quantile), respectively. To consider a wider range of precipitation regime, we multiple the daily mean precipitation in the wet year of 1972 by a factor of 2.25, 2.0, 1.75, 1.5, and 1.25 two create five very wet scenarios (i.e., highest annual precipitation = 1,669.8 x 2.25 = 3,757.1 mm yr^-1^). We also expanded the degree of dry conditions by divided the precipitation in the very year of 2007 by 2 (i.e., lowest annual precipitation = 528.6/2 = 264.3 mm yr^-1^). In total, we created 10 precipitation scenarios, with annual precipitation ranging from 264.3 to 3,757.1 mm yr^-1^. These precipitation scenarios were imposed on the model to investigate its effect on the presence/absence of regular patterns and the time it required for patterns to form.

To test the effect of soil clay content on pattern formation, we used the observed clay profile as the baseline (Fig. S6D). We multiplied a constant to this clay content vertical distribution to create new scenarios. The constant ranges from 0.1 to 2.2 with an increment of 0.14. This creates a total of 16 scenarios of clay profiles, with the mean clay content between the soil depth between 1.0 and 1.5 m (the likely pattern formation zone) varying from 4.45% to 97.9%.

