## Supplementary Materials for "The Primacy of Temporal Dynamics in Driving Spatial Self-organization of Soil Redox Patterns"

**Table S1**. Climatic and soil conditions of sites where regular iron redox patterns in soils have been reported. Note that we only collected papers that explicitly describe that the redox pattern is regular or show images of regular patterns.

| Location | Calhoun, SC, U.S. (Fimmen et al. 2008) | Santa Cruz, CA, U.S. (Schulz et al. 2016) | Southwest Victoria, Australia (MacEwan, Dahlhaus, and Fawcett 2012) |
|---|---|---|---|
| Photo of the pattern | 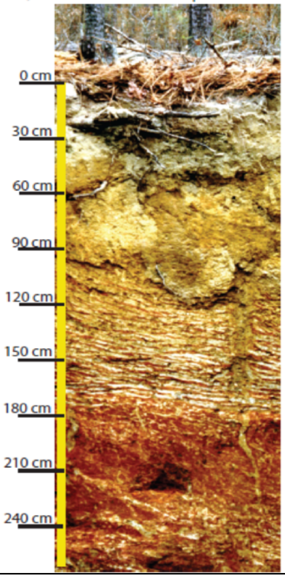  | 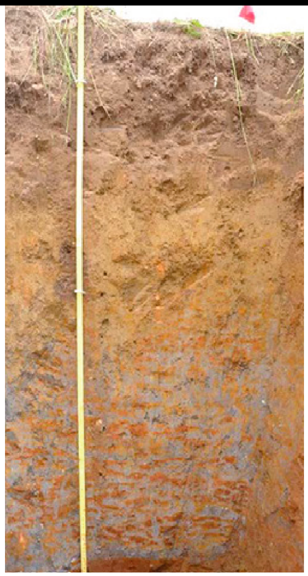 | 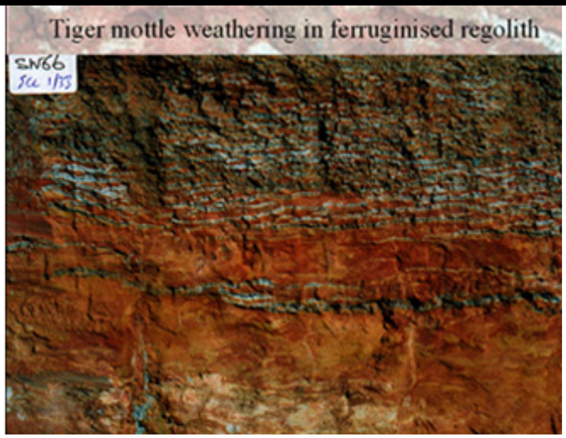  |
| Pattern formation soil depth | 1.2-1.8 m depth | 0.8-1.35 m depth | 1.8-2.1 m depth |
| Width | Orange: 1.28 cm; Grey: 1.92 cm | Orange: 2.8 cm; Grey: 2.8 cm | Orange: 2.0 cm; Grey: 1.0 cm |
| Mean annual temperature (°C) | 16.9 | 13.2 | 14.8 |
| Annual precipitation (mm $yr^{-1}$) | 1,270 | 730 | 1,092 |
| Clay content (orange layer) | 26.6% | 24% | – |
| Clay content (grey layer) | 73% | 56% | – |
| Background clay content | 44.5% | 51% | 48% |
| SOC (orange layer) | 0.08% | 0.062 – 0.19% | – |
| SOC (grey layer) | 0.2% | 0.17% – 0.28% | – |

**Table S2.** Comparison between the measured and modeled Fe(III) concentration, OM content, and clay content in the gray layers and in the orange layers of the formed pattern at our study site.

| | Grey (rhizosphere) microsites | | | Orange (iron-rich) microsites | | |
|---|---|---|---|---|---|---|
| | **Measured** | **Modeled** | **Relative error** | **Measured** | **Modeled** | **Relative error** |
| **Fe(III) (mol $m^{-3}$)** | 137.9 | 150.06 | +8.8% | 961.9 | 1368.94 | +42.3% |
| **OM (%)** | 0.50 | 0.51 | +1.9% | 0.203 | 0.149 | -26.5% |
| **Clay (%)** | 73 | 70.67 | -3.2% | 26.6 | 18.4 | -30.8% |

**Table S3.** The major hypothesized mechanisms, their corresponding model representations, and differences in their model predictions of redox patterns. The terms that are modified for each hypothesis are highlighted in gray.

| Mechanisms | Modifications to the Baseline Model | Distinguish among alternative mechanisms |
|---|---|---|
| Amplifying SDF + threshold dependent SDF (*Baseline model*)<br>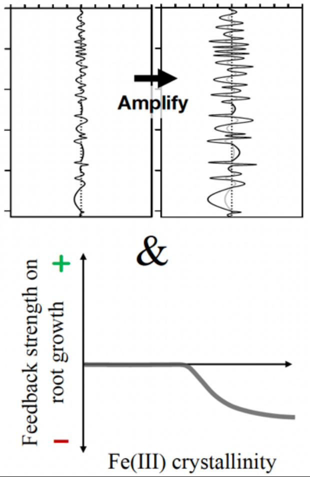<br> | $R_{Fe(III)} = 4k_1[OM]\frac{[Fe(III)]}{[Fe(III)]+H_{sFe3}}\frac{H_{FeO2}}{[O_{2L}]+H_{sO2}}Se^{\varphi} \quad (Eq.4)$<br><br>$G_B = \left(k_0 + k_g\frac{[O_{2L}]}{[O_{2L}]+H_{BO2}}\right)[B]f_{Fe(III)} \quad (Eq.10)$<br><br>$f_{Fe(III)} = \begin{cases}1, & [Fe(III)] \le [Fe(III)]_{ng} \\ \frac{H_{BFe}^2}{H_{BFe}^2+([Fe(III)]-[Fe(III)]_{ng})^2}, & [Fe(III)] > [Fe(III)]_{ng}\end{cases} \quad (Eq.11)$ | • **Pattern:** Modeled patterns of both OM and Fe(III) match the observed pattern.<br>• **Time:** ~ 900 years required to form patterns<br>• **Assessment:** The most plausible mechanism<br>• Video 1 |
| Amplifying SDF *only*<br>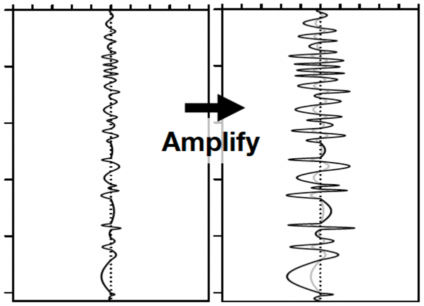<br> | $G_B = \left(k_0 + k_g\frac{[O_{2L}]}{[O_{2L}]+H_{BO2}}\right)[B]f_{Fe(III)} \quad (Eq.10)$<br>**** Set $f_{Fe(III)} = 1$ | • **Pattern:** Modeled patterns of OM and Fe(III) do not match observed pattern: (a) banding width much thinner; (b) layers of high (low) Fe(III) show high (low) OM matter, opposite of observations.<br>• **Time:** ~7,000 years required to form patterns<br>• **Assessment:** Unlikely to be the mechanism<br>• Video 2 |

| | | |
|---|---|---|
| Threshold-dependent SDF *only*<br>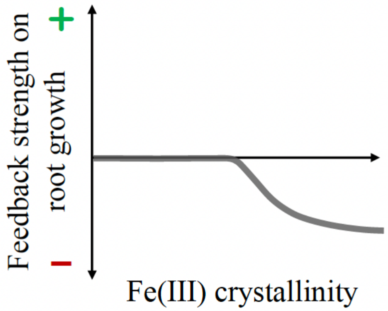 | $$R_{Fe(III)} = 4k_1[OM]\frac{[Fe(III)]}{[Fe(III)] + H_{sFe3}}\frac{H_{FeO2}}{[O_{2L}] + H_{sO2}}Se^{\varphi} \quad (Eq. 4)$$<br>**** Set $Se^{\varphi} = 1$ | • **Pattern:** No patterns form<br>• **Assessment:** Unlikely to be the mechanism<br>• Video 3 |
| Monotonic SDF *only*<br>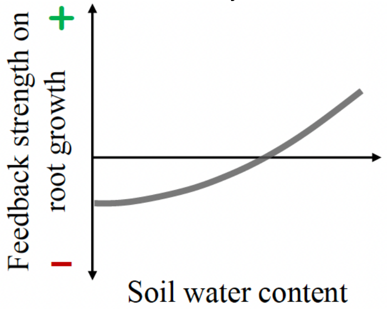 | $$R_{Fe(III)} = 4k_1[OM]\frac{[Fe(III)]}{[Fe(III)] + H_{sFe3}}\frac{H_{FeO2}}{[O_{2L}] + H_{sO2}}Se^{\varphi} \quad (Eq. 4)$$<br>**** Set $Se^{\varphi} = 1$<br>$$G_B = \left(k_0 + k_g\frac{[O_{2L}]}{[O_{2L}] + H_{BO2}}\right)[B]f_{Fe(III)} \quad (Eq. 10)$$<br>**** Replace $f_{Fe(III)}$ with $Se^2$ (Calabrese et al. 2020) | • **Pattern**: OM content much higher than the observation in gray layers is predicted by the model.<br>• **Time**: ~ 900 years required to form patterns<br>• **Assessment**: Not likely for our study site, but likely in dry environments and/or sandy soils<br>• Video 4 |
| Root template effect<br>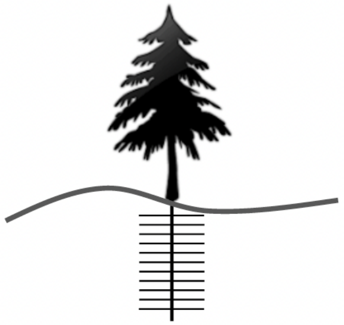 | $D_b = 0$ (*Eq.* 7)<br>$$R_{Fe(III)} = 4k_1[OM]\frac{[Fe(III)]}{[Fe(III)] + H_{sFe3}}\frac{H_{FeO2}}{[O_{2L}] + H_{sO2}}Se^{\varphi} \quad (Eq. 4)$$<br>**** Set $Se^{\varphi} = 1$<br>$$G_B = \left(k_0 + k_g\frac{[O_{2L}]}{[O_{2L}] + H_{BO2}}\right)[B]f_{Fe(III)} \quad (Eq. 10)$$<br>**** Set $f_{Fe(III)} = 1$<br>**** Set the regular root template in the initial condition (Fig. S2) and impose the same regular pattern on the root carrying capacity, $K_B$ (*Eq.* 12) | • **Pattern**: Modeled patterns of both OM and Fe(III) match the observed pattern.<br>• **Time**: ~ 200 years required to form patterns.<br>• **Assessment**: Unlikely to be the mechanism.<br>• Video 5 |

**Table S4**. Definition, notations, and values of model parameters.

| Parameters | Definition and unit | Values | References |
|---|---|---|---|
| $k_1$ | First-order Fe(III) reductive rate by organic matter ($d^{-1}$) | 0.000274 | (Chen et al., 2018) |
| $k_{app}$ | The pH-dependent Fe(II) oxidation rate constant ($d^{-1}$) | 21.6 | (Chen et al., 2018) (Chen et al., 2003) |
| $H_{sO2}$ | Half-saturation constant regulating the effect of dissolved oxygen concentration on Fe(III) reduction, Fe(II) oxidation, and root respiration (mol $m^{-3}$) | 0.02 | (Van Cappellen and Wang 1996) |
| $H_{sFe3}$ | Half-saturation constant regulating the effect of Fe(III) concentration on Fe(III) reduction (mol $m^{-3}$) | 100 | (Roden and Wetzel 2002) |
| $k_r$ | Root respiration rate constant ($d^{-1}$) | 8.64 x $10^{-5}$ | (Dong et al. 2018) |
| $k_d$ | Root decay rate constant ($d^{-1}$) | 0.0053 | Calibrated |
| $k_0$ | Root growth rate constant ($d^{-1}$) | 0.0053 | Calibrated |
| $k_g$ | Root growth rate constant affected by oxygen availability ($d^{-1}$) | 8.64 x $10^{-5}$ | Calibrated |
| $K_B$ | Carrying capacity of root biomass, which decreases exponentially with soil depth (g $m^{-3}$) | $1162e^{-0.849depth}$ | Fitted with field observations |
| $H_{BFe}$ | Half-saturation constant for the effect of Fe(III) on root growth (mol $m^{-3}$) | 200 | Calibrated |
| $[Fe(III)]_{ng}$ | Threshold Fe(III) concentration, above which Fe(III) imposes a negative effect on root growth (mol $m^{-3}$) | 650 | Calibrated |
| $M_{om}$ | Molar mass of organic matter (g $mol^{-1}$) | 30 | Represented by the generic form of $CH_2O$ |
| $D_{10}$ | Diffusion coefficient of $Fe^{2+}$ in soil water ($m^2$ $d^{-1}$) | 8.64 x $10^{-5}$ | (Kappler et al. 2005) |
| $D_{1L0}$ | Diffusion coefficient of dissolved $O_2$ in soil water ($m^2$ $d^{-1}$) | 1.9 x $10^{-4}$ | (Aachib, Mbonimpa, and Aubertin 2004) |
| $D_{1G}$ | Diffusion coefficient of gas $O_2$ ($m^2$ $d^{-1}$) | 1.754 | (Aachib, Mbonimpa, and Aubertin 2004) |
| $D_b$ | Diffusion coefficient of root biomass ($m^2$ $d^{-1}$) | 8.64 x $10^{-8}$ | Calibrated |
| $D_c$ | Diffusion coefficient of clay ($m^2$ $d^{-1}$) | 8.64 x $10^{-12}$ | Calibrated |
| $l$ | Pore connectivity parameter | 0.5 | (Mualem 1976) |
| $\beta$ | Parameter controlling magnitude of fluctuation in the initial condition of root biomass density | 0.5 | Derived from field measurements (Fig. S3) |
| $\varphi$ | Exponent controlling the effect size of normalized volumetric water content *Se* | 1.0 | Calibrated |

Video 1: Formation of redox patterns in upland soils by the mechanism coupling threshold-dependent SDF and amplifying SDF. The first row of plots shows profiles of the various chemical concentrations in the entire model domain (0-to-4-meter soil depth), including—from left to right—dissolved $O_2$ concentration (%), dissolved Fe(II) (mol $m^{-3}$), Fe(III) deposit (mol $m^{-3}$), and organic matter (mol $m^{-3}$). The second row of plots shows the pattern in the zone between 1.2 and 1.5 m soil depth. From left to right: soil volumetric water content, concentration of Fe(III) deposit (mol $m^{-3}$), clay content (%), and organic matter content (mol $m^{-3}$). The simulation shows a time interval of 1 year over ~ 800 years.

**Figure S1.** Model simulated patterns with five hypothesized mechanisms: Patterns formed by root template effect (A1-5); by coupling the amplifying SDF with threshold dependent SDF based on negative root responses to Fe(III) (B1-5); by the amplifying SDF alone (C1-5); by the threshold dependent SDF based on negative root responses to Fe(III) alone (D1-5); and by the monotonic SDF based on root responses to soil water content (E1-5). Blue lines in each plot show the observed background concentrations, represented by the data from nearby sites without patterning. Red points are empirical measurements from the Fe(III) concentrated orange layers at our study site, and the light-yellow points are empirical measurements from the Fe(III) depleted gray layers. The first column describes the modeled Fe(III); the second column is a close-up view of the Fe(III) in the pattern formation zone between 1.2 and 1.5 m soil depth; the third column describes the modeled organic matter (OM); the fourth column is a close-up view of OM in the pattern formation zone; and the last one is overlapping Fe(III) and OM profiles to show their spatial relationships of high and low.

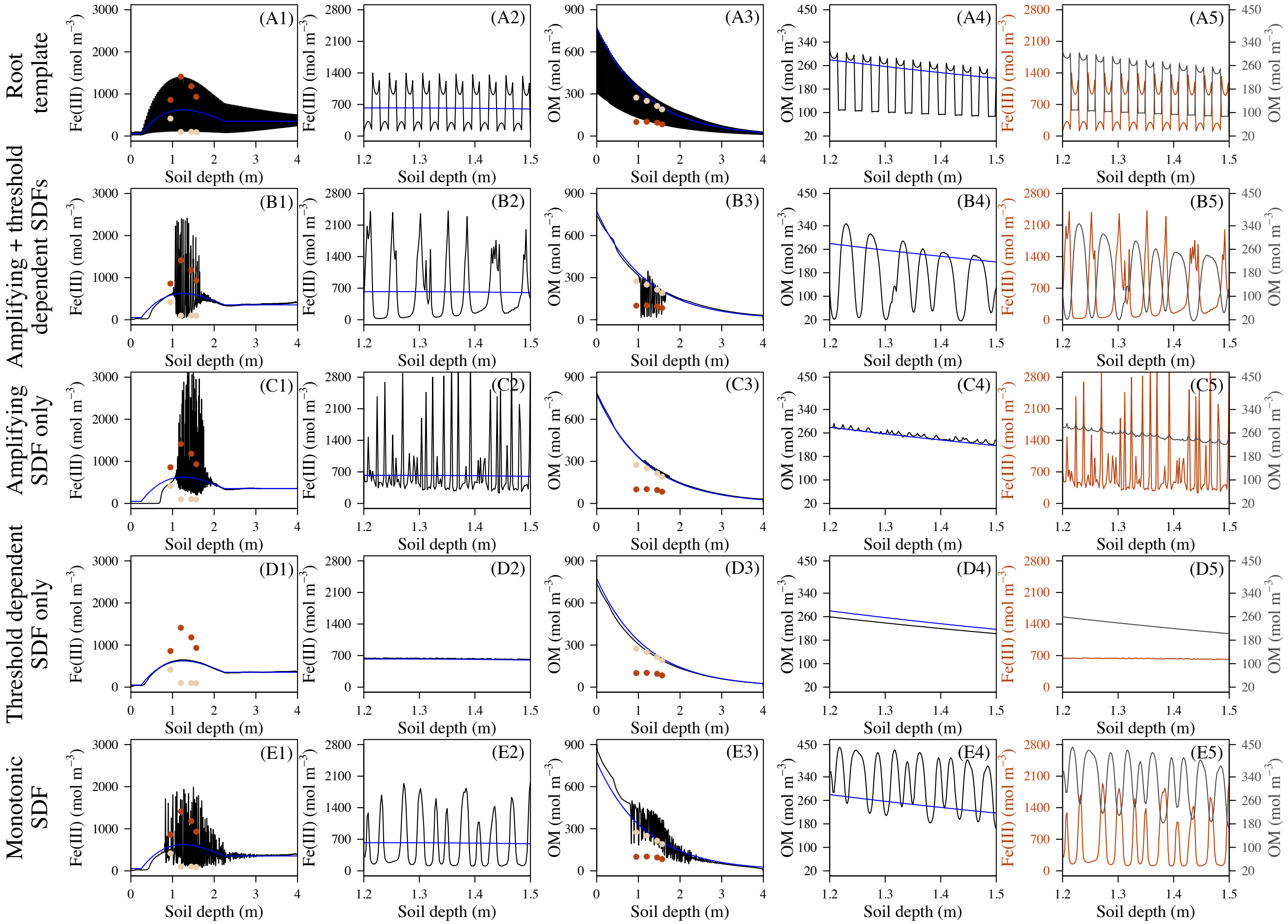

**Figure S2.** Redox patterns follow the *exact* pattern set by the root biomass in the initial condition. (A) shows the root biomass in the steady state (green) and in the initial condition (gray; as it overlaps exactly with steady-state distribution, gray line is not visible); (B) shows the Fe(III) profile in the steady state (orange) and in the initial condition (black); and (C) zooms in to the soil depth between 1.3 m and 1.5 m showing that root biomass in the initial condition (thick gray line) and in the steady state (green) and the initial-state (thick black line) and steady-state (orange) Fe(III) concentration distribution.

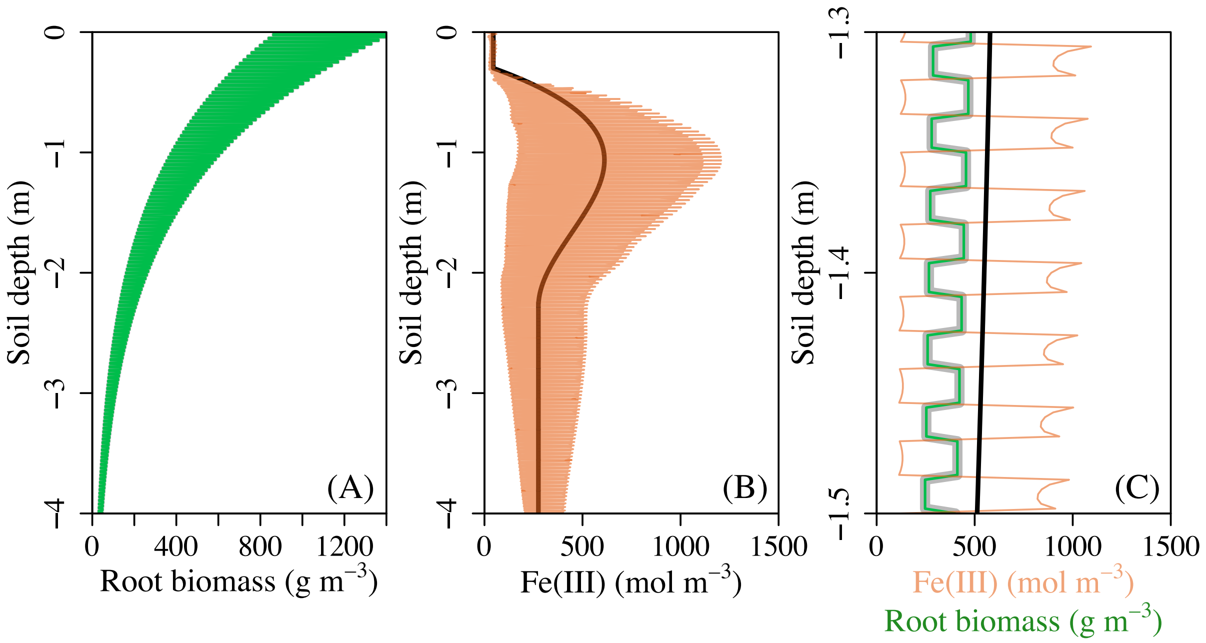

**Figure S3.** Difference of soil water content between orange and grey layers along the anoxic-oxic gradient, represented by the modeled soil water content in the pattern formation zone between 1.0 and 1.5 m soil depth. When soil is under anoxic condition, the difference of water soil content between orange and gray layers is small, resulting in weak amplifying effect. As soil gets drier and becomes more oxic, the soil water content between orange and gray layers becomes large, allowing a relatively strong amplifying effect occur. However, under oxic conditions, reductive dissolution of Fe(III) is suppressed. As a result, the amplifying effect is again limited. DO concentrations represent the average concentration between the soil depths of 1.0 and 1.5 m.

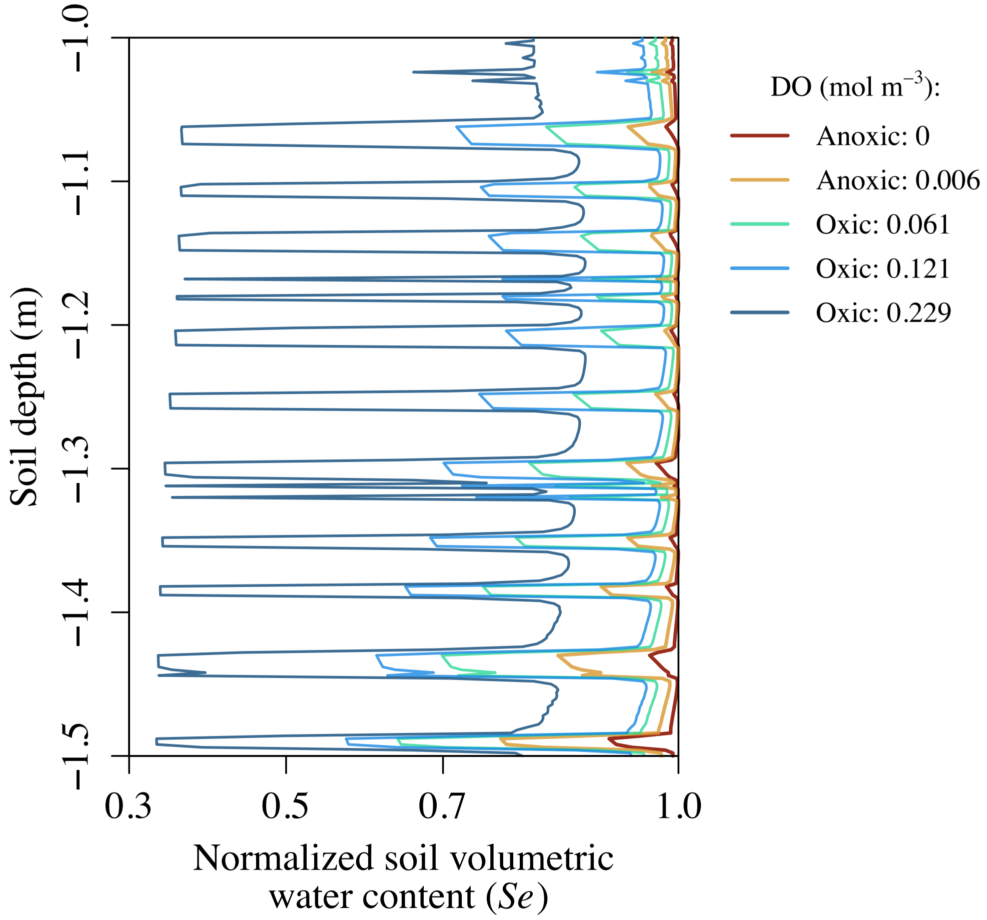

**Figure S4.** Comparison of the observed (between July 1, 2018 and July 1, 2019) and modeled $O_2$ concentrations at 50 cm (A) and 150 cm (B) soil depth and soil volumetric water content (C). Our model provides a good fit to the empirical data.

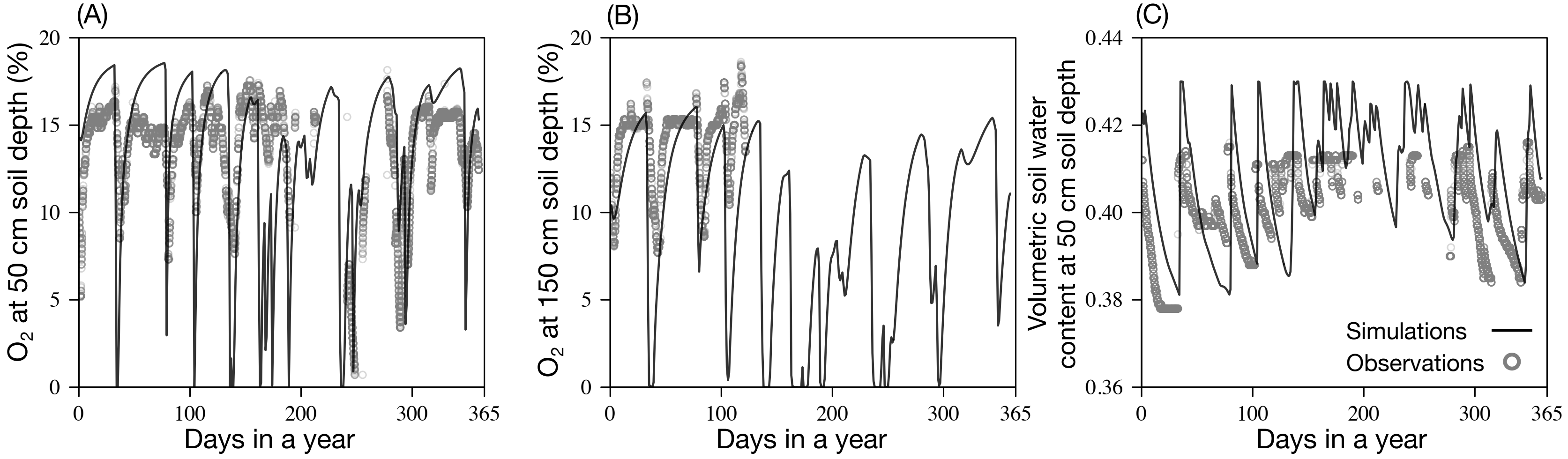

**Figure S5.** Modeled water retention curve and hydraulic conductivity curve for the soil texture in the background condition and for the contrasting soil textures in gray layers and orange microsites in regular redox patterns (Table S5 in Supplementary Text provides descriptions of characteristics of these three types of soil texture). Suction = 0 – pressure head, *h*.

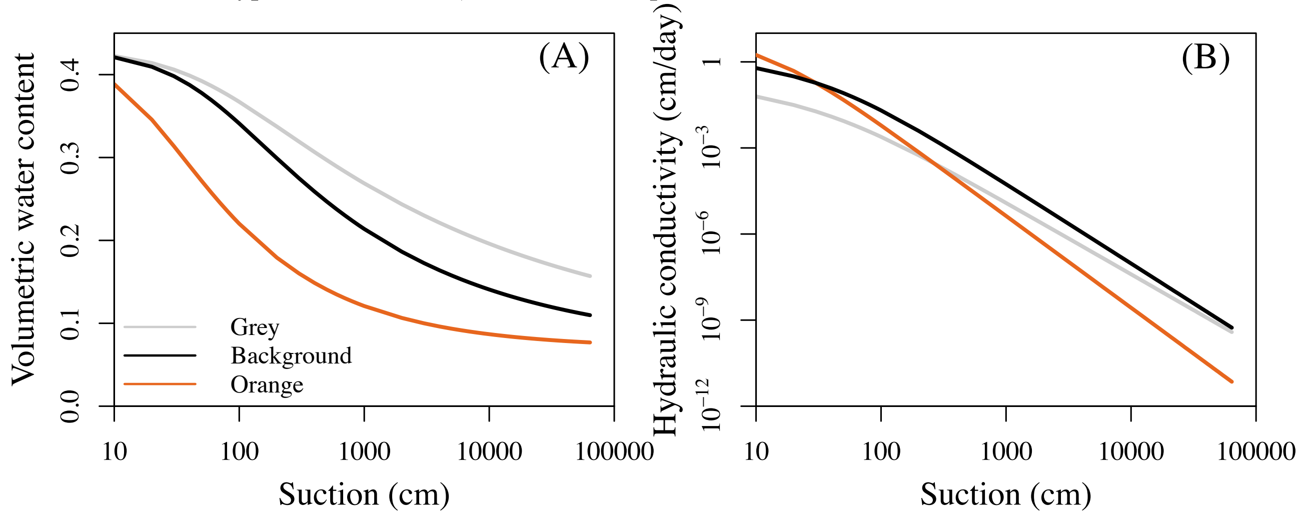

**Figure S6.** Empirical relationships between five hydraulic parameters (Table S5; Text S1) and clay content used in the model: (A) saturated hydraulic conductivity, $K_s$, (B) saturated soil water content, $\theta_s$, (C) residual soil water content, $\theta_r$, (D) fitting parameter, $\alpha$ (*Eq*. 19) (E) fitting parameter, *n* (*Eq*. 19). In (B)-(E): orange circles represent samples from orange layers of redox patterns at our study site, gray circles represent samples from gray layers, and black circles represent samples from bulk samples, as background level. Piecewise linear regressions were used to represent the relationship between the four hydraulic parameters and clay content (B-E), and a fitted exponential relationship, $K_s = 807241 \text{x} (\text{clay content})^{-3.24991}$, was used for the relationship between $K_{sat}$ and clay content (A). These empirical relationships were used to model soil water dynamics.

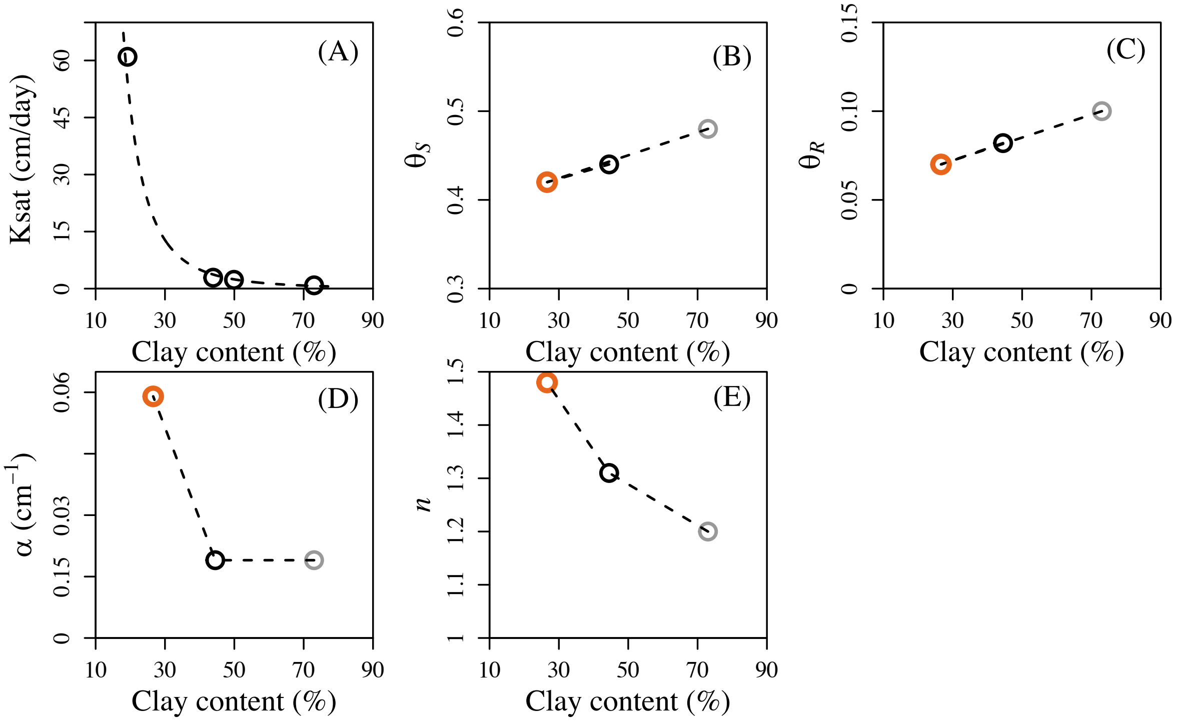

**Figure S7.** Initial conditions used in the model for (A) soil vertical profiles of root biomass, (B) organic matter concentration (OM), (C) Fe(III) concentration, and (D) clay content. The initial conditions are informed by observed patterns (red points) at nearby sites without patterns.

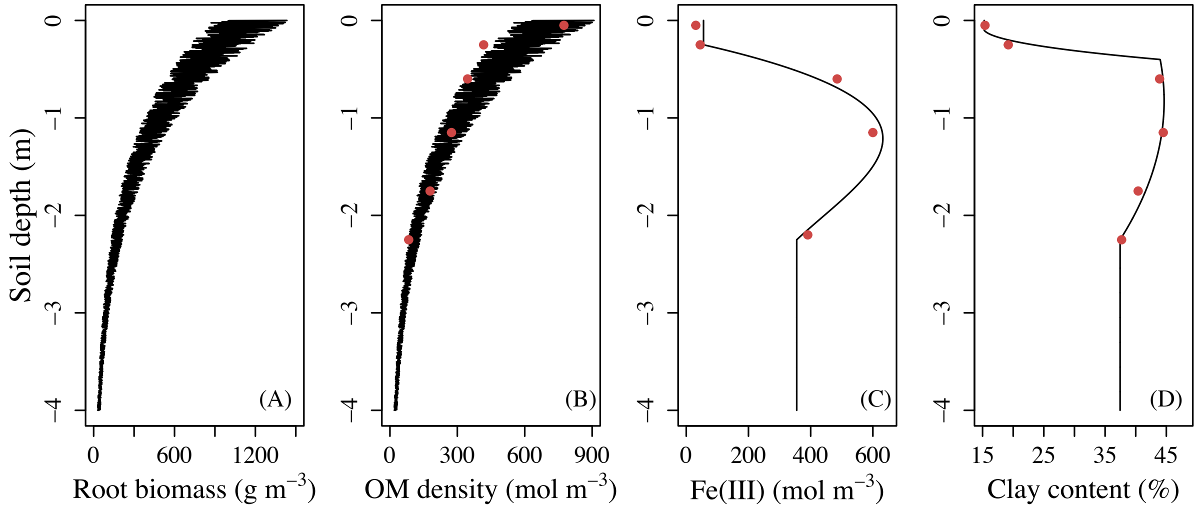

**Figure S8.** Sensitivity of Fe(III) reduction rate to soil water content ($\varphi$ in *Eq*. 4) affecting the spatial extent of pattern formation zone in the soil profile.

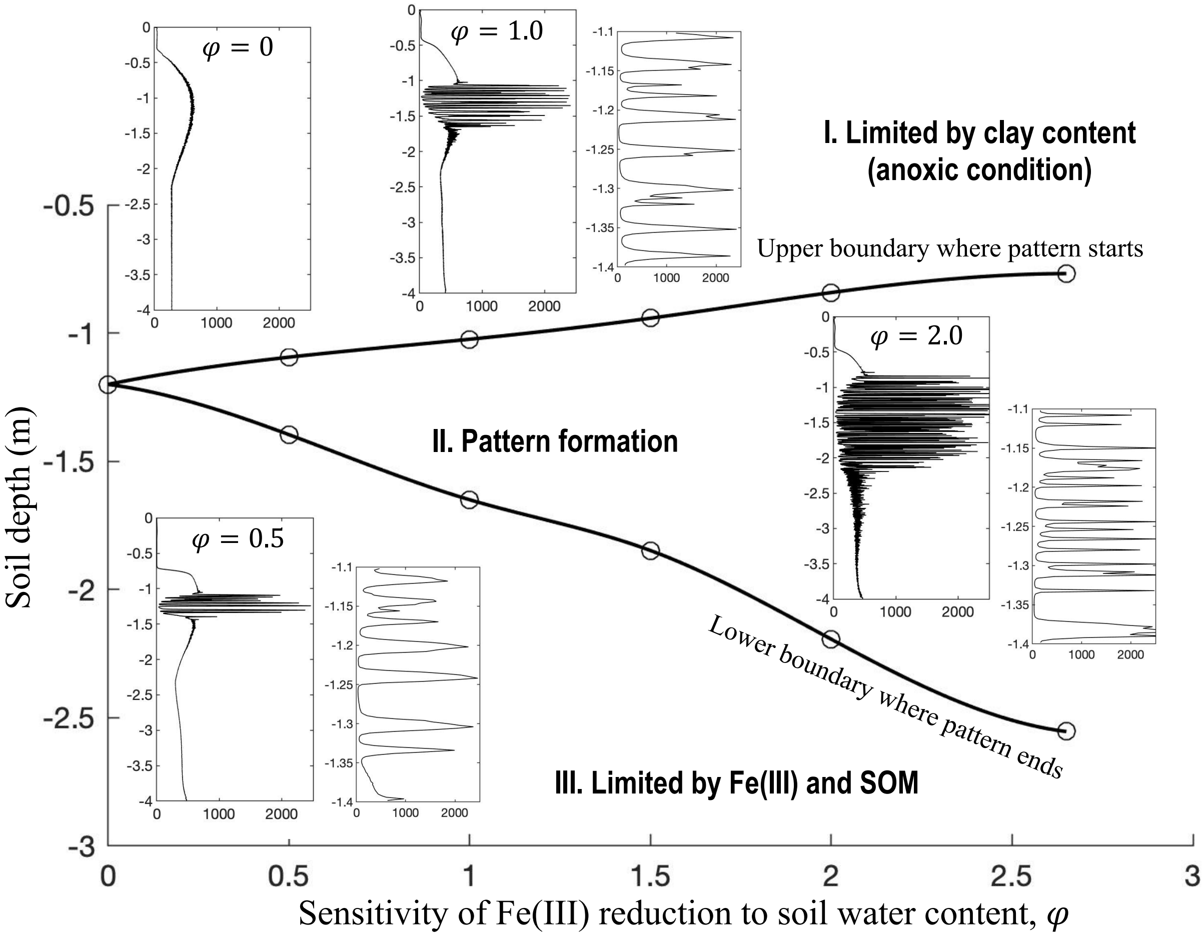

**Figure S9.** Time required for redox pattern formation in upland soils affected by first order Fe(III) reduction coefficient by soil organic matter, $k_1$ (A) and by annual precipitation (B). Pattern formation time changes linearly with $k_1$, but exponentially with precipitation ($R^2$ > 0.99).

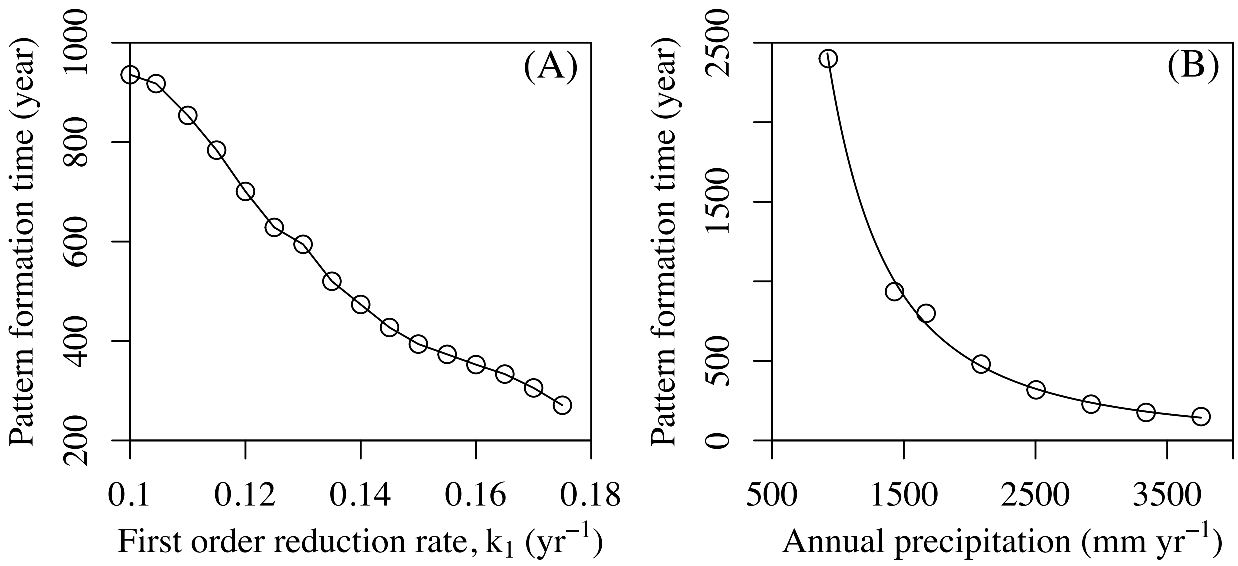

**Figure S10.** Measured (at every 1 cm) fine root distribution at ten different sites of the Calhoun Experimental Forest (South Carolina, U.S.) with best fit exponential decay functions. Dominant plant species varies among sites: hardwood trees ("*H*"), pine trees ("*P*"), and cottons ("*C*"). Biomass of cotton roots declines more rapidly than that of hardwood trees or pine trees. The dashed lines in each plot are 1.6 cm apart, the average width of an orange or gray layer in the regular redox patterns in our study site. Roots do not show regularly spaced (at 1.6 cm interval) biomass distribution (as in Figure S2).

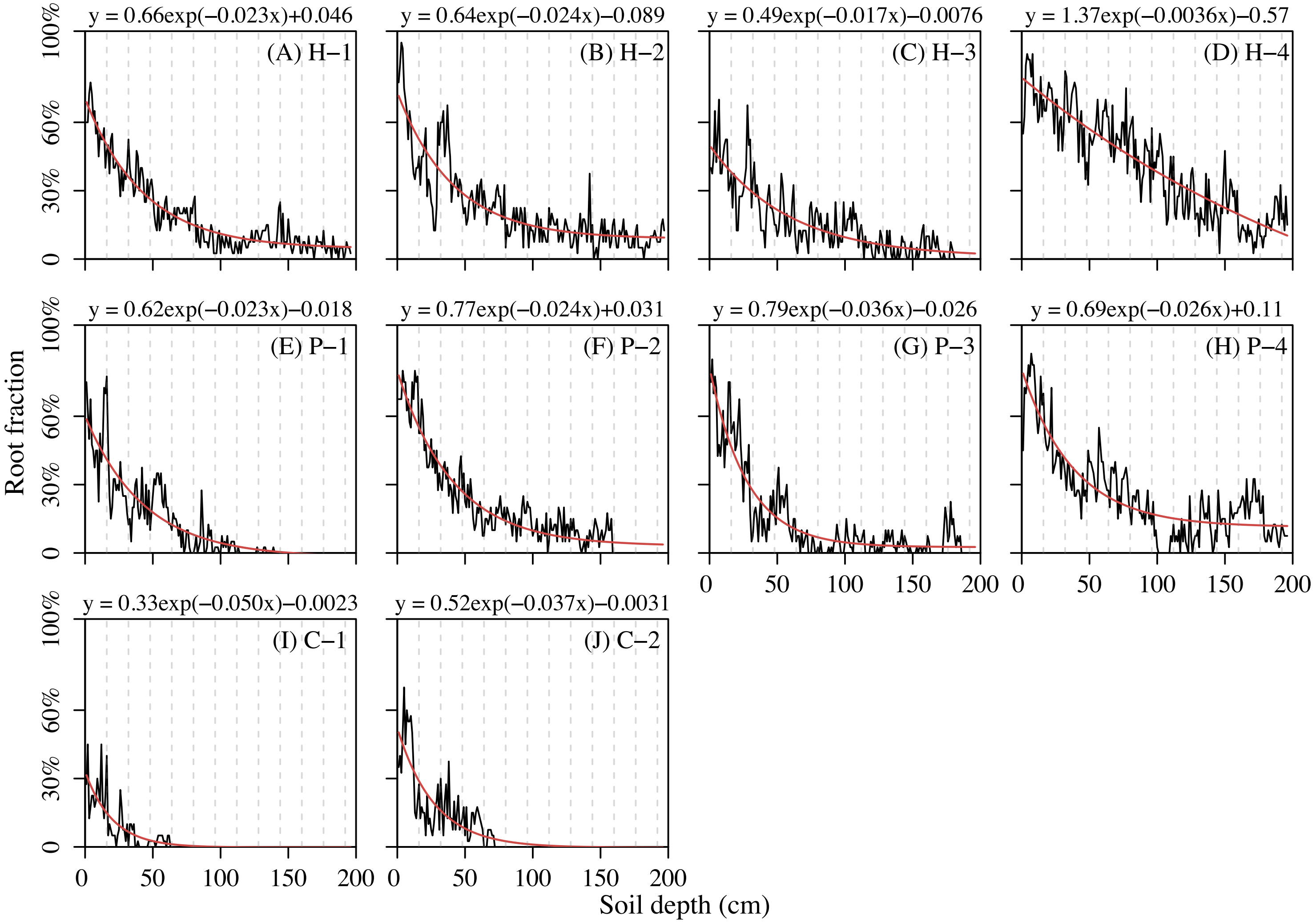

**Figure S11.** The effect of soil water content on the rate of Fe(III) reduction. The data are extracted from Figure 2 of the paper by Hodges et al. (2018). Fe(III) reduction rate represents the fraction of Fe(III) removed by reductive dissolution after two weeks of lab experiments, controlled at different levels of soil water saturation.

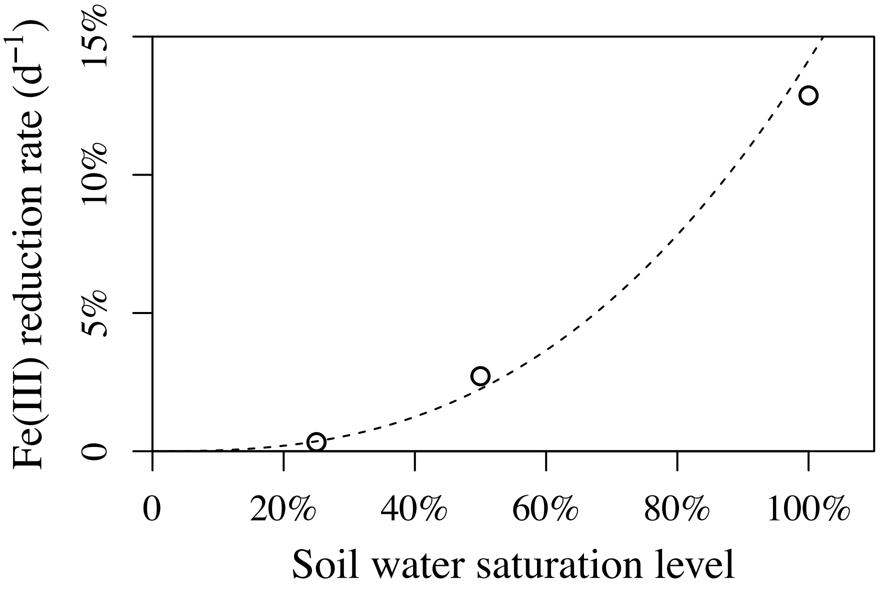

**Figure S12.** Empirical relationship between Fe(III) concentration and clay content with data from the Calhoun Experimental Forest (South Carolina, U.S.). This suggests the effects of iron redox reactions on clay disintegration and coagulation. Different colors represent the source of the data: Data from orange and gray layers in the regular redox patterns are in orange and gray circles, respectively; and data from sites without redox patterns, representing the background condition, are in black.

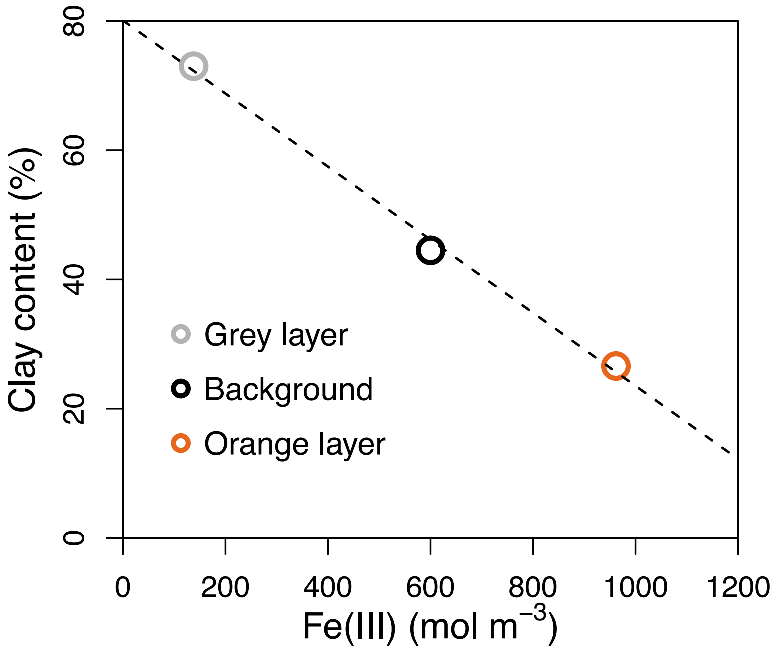

### Supplementary Text S1

#### Relationships between soil hydraulic parameters and clay content in soils

The progression of iron redox reactions continuously modifies clay content in soils, hence changing soil texture. Soil texture has a significant effect on water content and water flux, which in turn feeds back to affect soil redox dynamics and iron chemical reactions. To capture such feedbacks, we modeled the dynamics of soil hydraulic parameters based on empirical measurements in our study site and model calibration. We considered the impact of clay concentration on the five hydraulic parameters in *Eqs.* 19-21: saturated hydraulic conductivity ($K_S$), fitting parameters $\alpha$ and *n*, saturated water content ($\theta_S$), and residual water content ($\theta_r$). As clay concentration increases, soil texture becomes finer, water permeability decreases, i.e., lower $K_S$, $\alpha$ and *n*, and water retention capacity increases, i.e., higher $\theta_S$ and $\theta_r$ (Simunek, Van Genuchten, and Sejna 2005; Brogowski, Kwasowski, and Madyniak 2014). To represent these dynamics in the model, we compiled the soil texture data measured at gray layers and orange layers in the pattern formation zone in our study site. Since it is not feasible to have reliable information on soil texture in the initial condition before the regular redox patterns were formed, we used soil texture data collected at a nearby site without regular patterns (defined as "background" level in this study) to represent the soil texture in the initial condition. The soil texture data and hydraulic parameters were extracted from (Fimmen et al. 2008) and shown in Table S5.

**Table S5.** Soil texture and hydraulic parameters for the grey and the orange layers and proxy of the initial condition with data collected from a nearby site without redox patterning ("*background*").

| Sites | Clay (%) | Silt (%) | Sand (%) | Soil texture | $\theta_s$ | $\theta_r$ | $\alpha$ (1/cm) | *n* |
|---|---|---|---|---|---|---|---|---|
| Gray | 73 | 10.3 | 16.6 | clay | 0.48 | 0.10 | 0.019 | 1.2 |
| Orange | 26.6 | 20.3 | 53.1 | Sandy clay loam | 0.42 | 0.07 | 0.059 | 1.48 |
| Background | 44.5 | - | - | clay | 0.44 | 0.082 | 0.019 | 1.31 |

.

Roden, Eric E, and Robert G Wetzel. 2002. "Kinetics of Microbial Fe (III) Oxide Reduction in

Freshwater Wetland Sediments." *Limnology and Oceanography* 47 (1): 198–211.

Ryan, Michael G, Robert M Hubbard, Silvia Pongracic, R J Raison, and Ross E M C Murtrie. 1996. "Foliage, Fine-Root, Woody-Tissue and Stand Respiration in Pinus Radiata in Relation to Nitrogen Status." *Tree Physiology* 16: 333–43.

Schaap, Marcel G., Feike J. Leij, and Martinus Th Van Genuchten. 2001. "Rosetta: A Computer Program for Estimating Soil Hydraulic Parameters with Hierarchical Pedotransfer Functions." *Journal of Hydrology* 251 (3–4): 163–76. https://doi.org/10.1016/S0022-1694(01)00466-8.

Schulz, Marjorie, Dave Stonestrom, Corey Lawrence, Tom Bullen, John Fitzpatrick, Emily Kyker-Snowman, Jane Manning, and Meagan Mnich. 2016. "Structured Heterogeneity in a Marine Terrace Chronosequence: Upland Mottling." *Vadose Zone Journal* 15 (2): 1–14. https://doi.org/10.2136/vzj2015.07.0102.

Simunek, Jirka, M. Th Van Genuchten, and M. Sejna. 2005. "The HYDRUS-1D Software Package for Simulating the One-Dimensional Movement of Water, Heat, and Multiple Solutes in Variably-Saturated Media." *University of California-Riverside Research Reports* 3: 1–240.

Vogel, T., M. Cislerova, and J. W. Hopmans. 1991. "Porous Media With Linearly Variable Hydraulic Properties." *Water Resources Research* 27 (10): 2735–41. https://doi.org/10.1029/91WR01676.
